# Role of Gut Microbiome in Neoadjuvant Chemotherapy Response in Urothelial Carcinoma: A Multi-Institutional Prospective Cohort Evaluation

**DOI:** 10.1101/2023.01.21.525021

**Authors:** Laura Bukavina, Rashida Ginwala, Mohit Sindhani, Megan Prunty, Daniel Geynisman, Ghatalia Pooja, Henkel Valentine, Adam Calaway, Jason R. Brown, Andres Correa, Kirtishri Mishra, Raymond Pominville, Elizabeth Plimack, Alexander Kutikov, Mahmoud Ghannoum, Mohammed ElShaer, Mauricio Retuerto, Robert Uzzo, Lee Ponsky, Philip H. Abbosh

## Abstract

Treatment with neoadjuvant chemotherapy (NAC) in muscle invasive bladder cancer (MIBC) is associated with clinical benefit in urothelial carcinoma. While extensive research evaluating role of tumor mutational expression profiles and clinicopathologic factors into chemoresponse has been published, the role of gut microbiome (GM) in bladder cancer in chemoresponse has not been thoroughly evaluated. A working knowledge of the microbiome and its effect on all forms of cancer therapy in BC is critical. Here we examine gut microbiome of bladder cancer patients undergoing NAC. Overall, there was no significant difference in alpha and beta diversity by responder status. However, analysis of fecal microbiome samples showed that a higher abundance of Bacteroides within both institutional cohorts during NAC was associated with residual disease at the time of radical cystectomy regardless of chemotherapy regimen. Group community analysis revealed presence of favorable microbial subtypes in complete responders. Finally, fecal microbial composition outperformed clinical variables in prediction of complete response (AUC 0.88 vs AUC 0.50), however, no single microbial species could be regarded as a fully consistent biomarker. Microbiome-based community signature as compared to single microbial species is more likely to be associated as the link between bacterial composition and NAC response.

## Introduction

Treatment with neoadjuvant chemotherapy (NAC) in muscle invasive bladder cancer (MIBC) is associated with clinical benefit in urothelial carcinoma. Cisplatin-based NAC prior to extirpative surgery confers a 6-8% overall survival (OS) benefit compared to surgery alone [1]. Historically, the two most commonly utilized regimens include dose-dense methotrexate, vinblastine, doxorubicin, and cisplatin (dd-MVAC) or gemcitabine and cisplatin (GC), with pathological complete response (pT0) after NAC at the time of surgery being widely adopted as a surrogate end point for improved overall survival. [2] Conversely, patients with residual disease or nodal involvement are at a high risk of recurrence and progression of disease.

Historically, clinical stage is often used to guide treatment, although clinicopathological features alone with molecular subtyping, such as presence of DNA damage repair genes (*FANCC, ATM*, and *RB1*), has offered limited prediction due to overall low prevalence (<15% of patients.) [3, 4] While extensive research evaluating role of tumor mutational expression profiles and clinicopathologic factors into chemoresponse has been published, the role of gut microbiome (GM) in bladder cancer in chemoresponse has not been thoroughly evaluated. Indeed, the GM is increasingly seen as important and modifiable aspect of anticancer therapeutic response.[5, 6]We hypothesized that gut microbiome composition has a positive influence on cancer therapy in bladder cancer. To test this hypothesis, we prospectively collected fecal microbiome samples from patients with MIBC starting treatment with either GC or ddMVAC across two American institutions. In addition, samples from partners living with patients during NAC were collected to identify changes caused by chemotherapy vs confounding environmental effects. Furthermore, to investigate the change in GM during carcinogenesis, using N-butyl-N-(4-hydroxybutyl)-nitrosamine (BBN) murine model, stool pellets were collected for microbiome profiling throughout the BBN exposure period and at the time of sacrifice. This research provides the largest assessment of gut microbiome and its association with response to NAC and facilitates specific microbial special investigation and functions associated with response.

## Methodology

### Patient cohort and Sample Collection

The study was conducted in accordance with recognized ethical guidelines and approved by Case Western Reserve/University Hospitals Cleveland Medical Center and Fox Chase Cancer Center under the IRB# STUDY20200350 and IRB #18-4001, respectively. Patients with MIBC undergoing cystectomy were enrolled prospectively across both institutions.

#### Case Western cohort

stool samples prior to cystectomy at CW starting in 2018 until 2021. Patients were excluded from participation if they have any antibiotics within 6 weeks of collection, or history of *Clostridium difficile* infection and treatment within 2 months. Pre-surgical stool samples were collected at the time of cystectomy prior to initiation of antibiotics with rectal swabs, and all collections were performed prior to the use of preoperative antibiotics for surgical preparation. The swabs were immediately placed into 1.5 ml microcentrifuge tubes containing 1 ml of phosphate-buffered saline (PBS). Swabs were stored at 0°C during transport to the laboratory for processing within 2 hours of collection. Samples were then resuspended and stored in sterile PBS at -80°C until analysis.

#### Fox Chase cohort

stool samples were collected prior to NAC (ddMVAC), after cycle two (of three), and prior to cystectomy. Stool samples from partners were collected at the same time and stored. Samples were collected at home using the GenoTek stool sample collection kit which contains a preservative. Samples were stored at room temperature for <24h prior to being split into aliquots and stored at 80°C until use. All patients and partners were provided an OMNIgene GUT kit (OMR-200) (DNA Genotek, Ottawa, Canada) for outpatient fecal sample collection, which maintains DNA stability at room temperate for up to 60 days.

The final analysis consisted of 29 bladder cancer patients and 26 non cancer healthy controls (CW), and additional 7 patients at FCCC with 7 matched partners.

### Murine studies

The C57BL/6 female and male mice were purchased from Taconic Laboratory. The animal experiments were carried out following Fox Chase Cancer Center Institutional Animal Care and Use Committee guideline (IACUC 19-03). BBN was purchased as a single batch from TCI America (Portland, Oregon, USA, Batch ODW3F-FH). Ad libitum BBN was administered at a concentration of 0.05% in water and replenished weekly to half of the cages, with the other half receiving regular drinking water from the same source. Half of the mice in each treatment arm were male. Mice began BBN administration via drinking water at 8-10 weeks of age and continued for 12 weeks. After 12 weeks of BBN exposure, mice were given regular drinking water until the end of the study. Presence of tumors was evaluated via excretory μCT urography following retroorbital injection of dilute Visipaque contrast. (Supplementary Figure 1) Stool pellets were collected at pretreatment, six weeks, twelve weeks and between 16 to 22 weeks (when tumors were visible as filling defects on μCT scan). A total of 23 mice were exposed to BBN (14 males, 9 females), with 18 water only controls (10 males, 8 females). When sacrificing mice, one mouse from each cage was sacrificed at each time point rather than sacrificing all mice from one cage to avoid confounding factors that might be present in one cage and not another. Stool pellets were collected from each mouse if it spontaneously defecated or with gentle abdominal pressure. Pellets were resuspended in PBS and stored -80□°C until use.

### DNA extraction and bacterial 16S rRNA sequencing

Fox Chase samples: DNA extraction from patient or murine stool samples was performed using the QIAgen kit.). Fecal samples at FCCC were stored in 250μl aliquots at -80°C for DNA extraction. DNA was extracted from the thawed fecal samples using the DNeasy blood and tissue kit (Qiagen). Briefly, to 250μl fecal sample, 180 μl Buffer ATL was added and mixed carefully. To this mixture 59 μl of enzyme cocktail containing 50 μl lysozyme (10 mg/ml, Sigma-Aldrich), 6 μl mutanolysin (25 KU/ml, Sigma-Aldrich), and 3 μl lysostaphin (4000 U/ml, Sigma-Aldrich) was mixed and the resuspended sample was transferred in to tubes containing 0.4g sterile zirconia beads. The samples were homogenized in a Mini-BeadBeater™ at maximum speed for 1 min and then incubated at 37°C for 30 min. 20 μl of proteinase K was added and thoroughly mixed followed by incubation at 56°C for 1-2h to allow complete lysis. The samples were then centrifuged at maximum speed for 5 min to pellet debri. The supernatant was then used for DNA extraction using the DNeasy blood and tissue kit according to manufacturer’s guidelines. Sample preparation and sequencing was performed in collaboration with Case Western Microbiome Core and Fox Chase Cancer Center.

16S rRNA gene sequencing methods were adapted from the methods developed for the NIH-Human Microbiome Project [7]. Briefly, the 16S rDNA V3-V4 region was amplified by PCR using primers with overhangs to facilitate addition of UAS, then indexed with primers containing the Illumina USA. Note that the first stage PCR used primers that include a 1-, 2-, 3-, or 4-basepair frameshift outside the target site on each side to eliminate the need for phi-X DNA on the sequencer as describe by Lundberg. [8] Samples were pooled at equimolar proportions sequenced on the MiSeq platform (Illumina, Inc, San Diego, CA) using V3 chemistry (2×300 bp paired-end kit). The V3-V4 regions provide adequate information for the taxonomic classification of microbial communities from specimens associated with humans. [9, 10]

Filtered sequences with >97% identity was clustered into Operational Taxonomic Units (OTUs), and classified at the genus against the SILVA 16S ribosomal RNA sequence database (release 138.1) [11]. The relative abundance of each OTU was determined for all samples. A step-by-step description of our analysis pipeline has previously been published and is available [9]. QIIME 2 platform was utilized for processing, and visualizing microbiome data. [12] All fastq files spreadsheet are available as supplementary index 1.

### Gut microbiome analysis

Complete response (CR) was defined as pT0/pTis N0 M0 pathological assessment, with all other stages defined as NR. Differentially abundant OTUs were identified using LEfSe [linear discriminant analysis (LDA) effect size] for each pairwise comparison of clinical groups (healthy vs. bladder cancer, bladder cancer vs partner, complete response (CR) vs no response (NR)). LEfSe first uses a nonparametric factorial Kruskal–Wallis sum-rank test to identify differentially abundant OTUs.[13] This is followed by a set of pairwise tests among clinical groups to ensure biologic consistency using the Wilcoxon rank-sum test. LDA is then used to estimate the effect size of each differentially abundant OTU. LEfSe statistics were ranked to identify the greatest differences in microbial relative abundance across patient groups mentioned above.

To create a genus-level OTU table, OTUs with the same genus name were merged into one genus. We calculated the relative abundance of each genus. Samples within bladder cancer patients were grouped into two clusters based on beta diversity using k-means clustering.[14] The number of clusters (*k* = 3) was determined heuristically.

### Statistical assessment

Alpha diversity comparison between bladder cancer vs healthy, CR vs NR, bladder cancer vs partner and enterotype were compared using Wilcoxon rank-sum or Mann-Whitney (MW) test (used for comparison between binary variables) and Spearman rank was used to compare continuous variables. Fisher’s exact test was used when proportions were compared between binary variables. Adjustments for multiple comparisons were done using the false-discovery rate (FDR) method at an α level of 0.05. Volcano plots were generated for log_10_(FDR-adjusted *p*-values) on the y-axis and median-adjusted effect sizes on the x-axis. Hypothesis testing was done using both one-sided and two-sided tests as appropriate at a 95% significance level. All analyses were conducted in R [15], and Python [16]. All visualization was performed via R, Python, with CorelDraw (™).

Logistic regression for response-prediction classifier using MicrobiomeMarker(1.3.3) utilized to test prediction performance of identified microbiome genera.[17] Using high dimensional microbiome data, the model creates a sub-community consisting of signature microbial taxa associated with the disease (BC) and area under the curve (AUC) evaluation. AUC was used to assess the performance of the classifier for complete response across both institutions independently and combined. Code for MicrobiomeMarker can be obtained <u>here</u>.

## Results

### Microbiota perturbations and bladder cancer specific alterations compared to healthy controls

We prospectively collected fecal microbiome samples from radical cystectomy candidates MIBC From two institutions (CW and FCCC) utilizing different NAC protocols (GC vs ddMVAC). Fecal samples were collected at time of surgery at CW and compared to healthy, non-cancer controls (CTR), while samples collected at FCCC were prospectively collected pre-, during-chemotherapy as well as at the time of surgery. In addition, samples of their partners living with the patients during treatment were collected at FCCC, to assess possible alterations due to environment, rather than disease or chemotherapy itself. Eligible patients at CW (n=28 BC, n=26 CTR), and FCCC (n=7 BC, n=7 partners).

We first assessed for GM differences in all available CW samples in BC as compared to CTR. BC patients had lower prevalence of *Escherichia-Shigella* (1.2% vs 9.6% p=0.001) and *Pseudomonas* (0.01% vs17.6%, p=0.001), with exaggerated abundance of *Prevotella* (5.5% vs 0.01%, p=0.01) in stool compared to CTR. There was overall increase in alpha diversity (Shannon p=0.00015, Simpson <0.00043 Chao p=0.0016, Observe p=0.00039) (Supplementary index 1) in the BC cohort although the differences were minimal compared to controls (Figure 1A, D, E). Weighted UniFrac distances using principal coordinate analysis (PCoA) representative for beta-diversity (Figure 1C) demonstrated a clear separation of communities (p<0.001) by presence of disease (BC vs CTR). Based on these data, we examined an enrichment through high dimensional class comparisons using LEfSe, which again demonstrated differentially abundant bacteria in the GM of healthy controls vs bladder cancer. In addition to confirming previously established increased abundance of *Pseudomonas* in healthy controls, we identified presence of *Fusobacterium* as significantly enriched in BC cohort (LDA>3.5, p<0.05). (Figure 1B, F)

**Figure 1:**
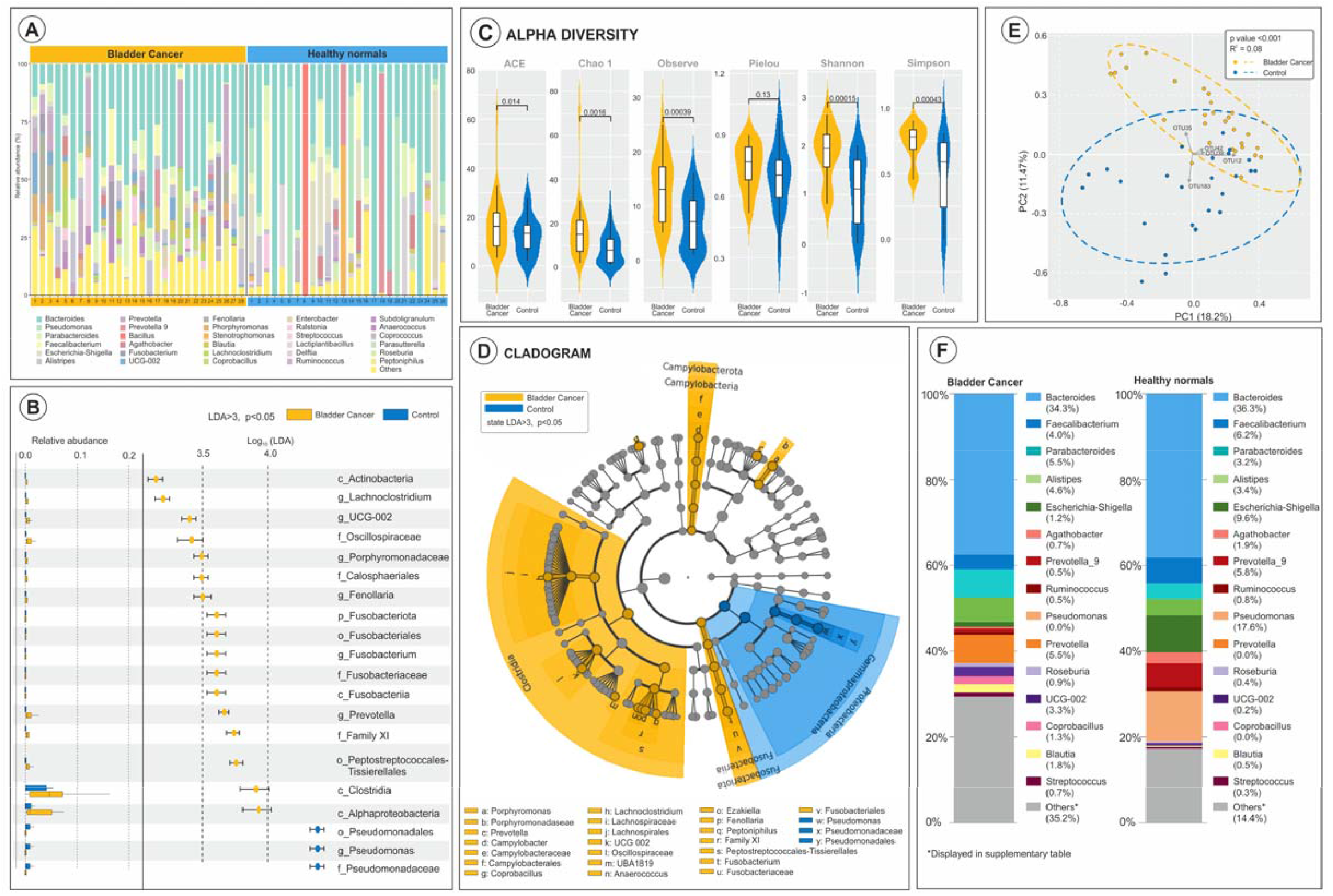
Case Western Bladder Cancer vs Controls microbiome diversity and differences. A) Stacked bar plot of phylogenetic composition of common bacteria taxa(>0.1%abundance) at the genus level in BC (n=28) and Controls(n=29) also visualized in (F). B) Differentially abundant bacteria (FDR adjusted) within all taxonomic levels up to genus. C) Alpha diversity scores of the gut microbiome in both cohorts, with error bars representing distribution of diversity scores. C) Beta Diversity depicted as Weighted UniFrac PCoA demonstrating separation of bacterial communities by disease status. D) Taxonomic cladogram from LEfSe showing difference in GM, across all taxonomic levels. Dot color representing group and composition.

Since the collection at CW cohort was taken at the time of surgery after chemotherapy, we then focused on defining gut compositional differences in BC patients prior to any intervention, during and after chemotherapy as compared to their partners. To test the influence of environment (diet, housing, exposure), we collected stool samples from live-in partners at the same timepoints. No major differences in alpha and beta diversity were noted throughout receipt of ddMVAC in patients (Figure 2 A, B, F), although minor fluctuations in the compositional differences were seen, particularly increasing prevalence of *Bacteroides, Escherichia-Shigella*, and *Roseburia* post NAC as compared to prior.(Figure 2 A, D) Overall, 8 OTUs were exclusive to the post-NAC samples (Figure 2E), however, overall diversity and distribution was similar throughout receipt of chemotherapy (102 OTUs common to all 3 timepoints). LDA analysis revealed no significant difference in GM composition between before, during or after chemotherapy (LDA>3, p=0.25) (Figure 2C).

**Figure 2.**
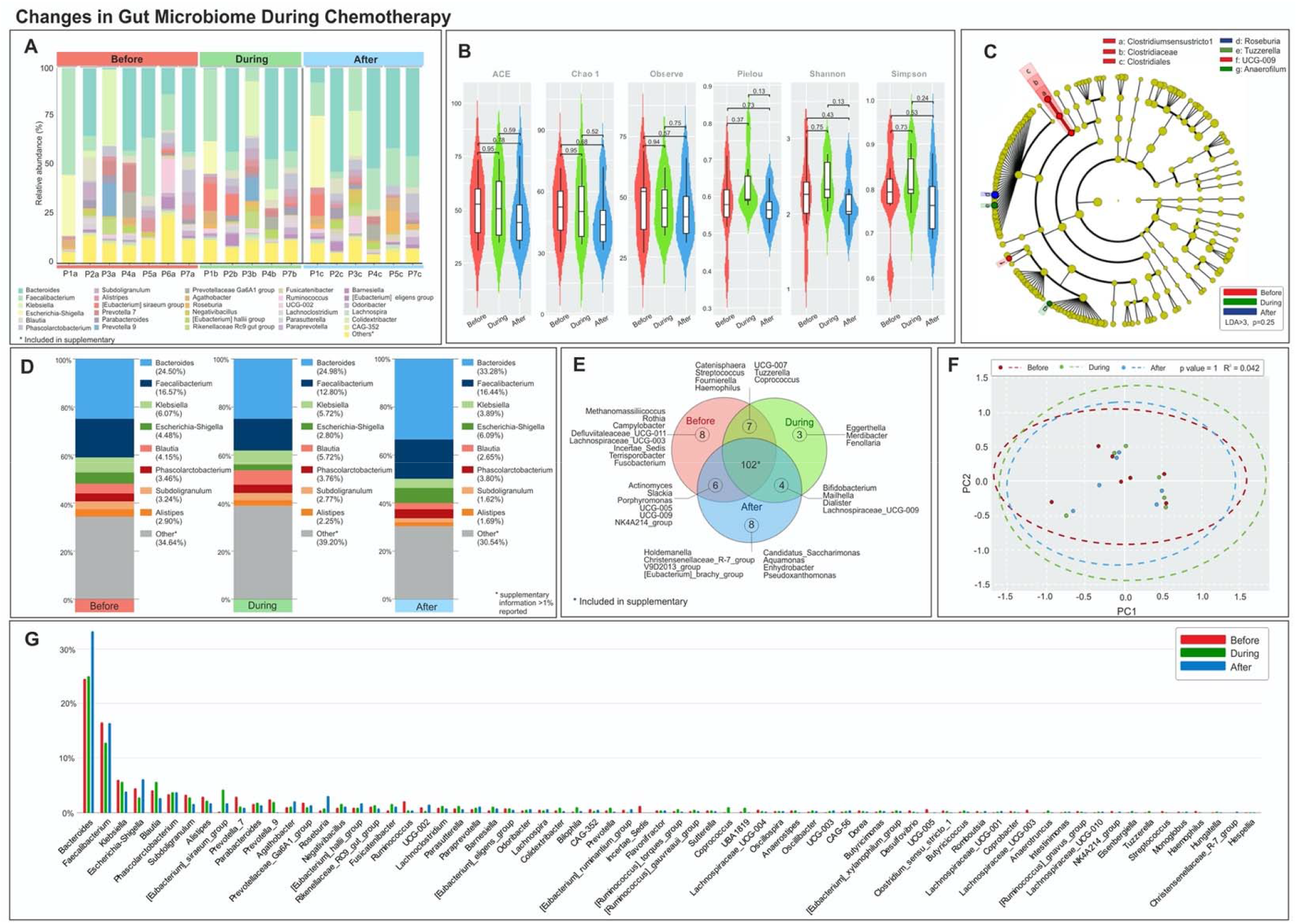
Changes in gut microbiome throughout neoadjuvant chemotherapy. A) Stacked bar plots of individual patients designated as letter and subsequent collection timing (1: before chemo, 2: during chemo, 3: after chemo) and composition of common bacteria taxa at genus level. (n=7). Additional phylogenic distribution is depicted in (D) and (G). Prevalence over 0.1% is reported in supplementary index 3. Limited differences in alpha diversity (B) and beta diversity (F) are visualized between treatment time points. (C) Taxonomic cladogram from LEfSe showing no statistically significant differences in GM during NAC cycle. E. Venn diagram denoting patients grouped by NAC time point, and shared/unique bacterial OTUs. Additional OTUs reported in supplementary index 3.

While both patients and partners did not exhibit any significant differences in alpha or beta diversity throughout NAC (Supplementary Figure 1), unlike BC patients who were noted to have an increase in *Bacteroides, Escherichia-Shigella and Roseburia* post chemotherapy, their partners’ abundance for *Bacteroides* decreased throughout the course of treatment (Supplementary Figure 1).

### Bacteria taxa impact on NAC treatment response

Since the compositional differences explored above may influence not only cancer development but similarly response to therapy, we wanted to next explore how specific gut microbiome composition impacts treatment response. To test this, we compared GM OTU abundance in CR vs NR cases across both institutions. Due to the differences in NAC regimen across both institutions, we performed individual institutional comparisons.

We first compared the enrichment of operational taxonomic units (OTUs) in CR vs NR (n= 4, and n=5, respectively) in the CW cohort where the standard NAC regimen is composed of gemcitabine and cisplatin (GC). As compared to CR, non-responders demonstrated distinct sets of abundant OTUs, as well as 10 unique OTUs. (Figure 3E) Specifically, *Bacteroides, Fusobacterium, Bifidobacterium, Anaerococcus* were enriched in the GM of non-responders (Figure 3 A, C), although no major differences in alpha or beta diversity were observed (Figure 3 B, D). Furthermore, high dimensional class comparisons using LEfSe (adjusting for overall abundance) demonstrated *Escherichia-Shigella* to be the *only* bacteria present to distinguish CR vs NR (LDA>3, p<0.01) (Figure 3H) while the remainder of rare low abundances were not confirmed. Presence of high *Escherichia-Shigella* (E^h^) prevalence as compared to low (E^l^) (20% vs 0%, p=0.054) associated with responder status, with none of the patients with E^h^ demonstrating CR at the time of surgery (Figure 3 F,G). Although not reaching significance, presence of *Bifidobacterium* was associated with CR (LDA>2, p<0.056) in CW cohort. (Figure 3H)

**Figure 3.**
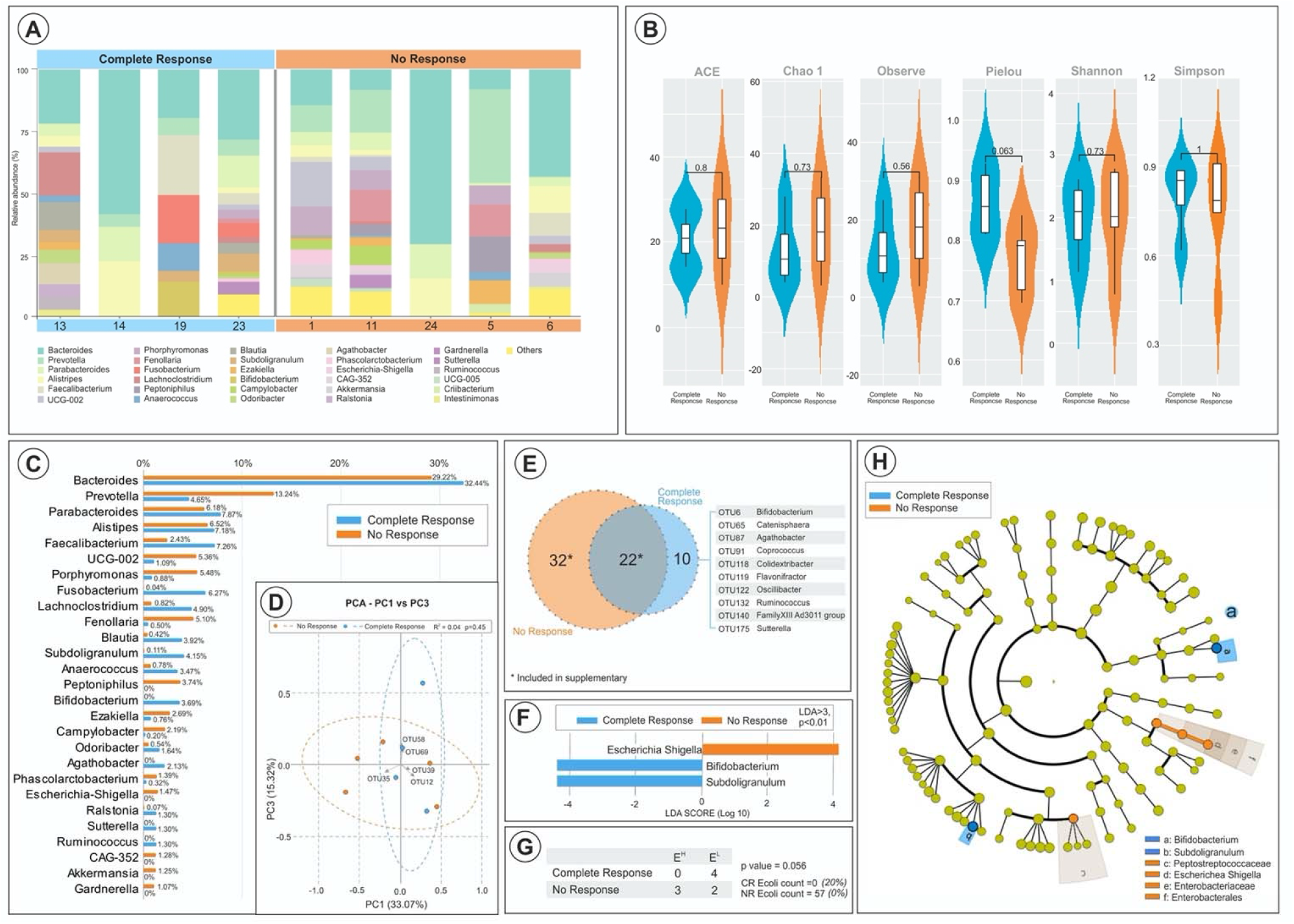
Responder status and compositional differences in GM in CW cohort. A. Columns denote patients grouped according to response status, and patient identification, with phylogenetic composition of common bacterial taxa. Direct comparison is also visualized in (C) based on composition of GM. (B) Alpha diversity scores of the GM in CR and NR to gemcitabine cisplatin NAC show minimal statistical difference. (D) Beta diversity via weighted UniFrac PCoA showing no evidence of clustering by group. (E) Venn diagram of GM based on responder status showing 10 unique bacteria OTUs present in CR, and 32 unique OTUs present in NR. (F) LDA scores computed for differentially abundant taxa in the GM of CR vs NR, with length of bar indicating the effect size calculated. (G) Composition of total E coli reads and percent prevalence average by CR vs NR. (H)Taxonomic cladogram from LEfSe showing differences in GM by NAC response status.

As the NAC regimen at FCCC is composed of ddMVAC, we explored the landscape of gut microbiome separately, noting enrichment in abundance of *Bacteroides (*34.49% vs 5.64%), *Klebsiella (7*.*07% vs 0%)* in NR vs CR, respectively. (Supplementary Figure 2 A, G) While no differences in alpha diversity (Supplementary Figure 2D) were observed, separate community clustering between CR vs NR was seen (p<0.001, Supplementary Figure 2B). This was again seen following LEfSe, with NR harboring increasing abundance of *Bacteroides* (LDA 4.9, p<0.05) compared to CR. (Supplementary Figure 2C, F). Within FCCC cohort, CR demonstrated 10 unique OTUs compared to NR (Supplementary Figure 2E).

Due to the district differences in composition and association of response based on NAC regimen, we asked whether *grouped abundances* within a specific treatment were related to CR, rather than single bacterial genus. We grouped all identified OTUs using hierarchical clustering without input of response data. We found that CW patients (BC and healthy controls) segregated into three community types. While group III contained only healthy controls (Supplementary Index 1), BC patients GM clustered were into two groups (I and II). (Supplementary Figure 3A). As expected, alpha and beta diversity differences were significant among the groups due to nature of the analysis. (Supplementary Figure 3 B, C) Group II consisted of non-responders only, with zero patients with Group II experiencing a complete response after GC therapy, while patients within Group I were more likely to experience CR (4/6, 66.67%). (Supplementary Figure 3E, F). As compared to Group 1, patients grouped within Group 2 had an overall increased abundance of *Bacteroides* (50.42% vs 16.01%), and *Pseudomonas* (0.2% vs 7.78%) with a decrease in *Prevotella* (0.71% vs 17.51%) (Supplementary Figure 3F). There were 40 unique OTUs within Group I (CR group) as compared to 7 unique OTUs in Group II (NR group). (Supplementary Figure 3D).

Within FCCC cohort (ddMVAC), similar community clustering into three community types was seen, with (3/4, 75%) *CR visualized in Group 1 only*, with majority of NR (11/14, 78.5%) segregating into group 2 (Supplementary Figure 4A, G). Similar to CW cohort, group II (with NR) was more likely to harbor enrichment of *Bacteroides* (41.22% vs 2.94%). (Supplementary Figure 4 A, E, G). Compared to CR, NR were more likely to experience a rise in *Bacteroides* throughout chemotherapy compared to CR (% △ + 15.05% vs -9.73%, p<0.001) ***Thus, higher abundance of Bacteroides within both CW and FCCC cohorts during NAC was associated with residual disease at the time of radical cystectomy regardless of chemotherapy regimen***.

### Establishment of response-predictor classifier

Using logistic regression, we constructed a response-prediction classifier in patients with MIBC and prediction for CR using a set of clinical variables as well as microbial genera. Our cohort consisted of 17 patients (27 stool samples) between two institutions (8 CR, 9 NR). Model selected 99 discriminatory genera to form a set of features to predict for NAC response based on machine learning modeling per institution. Following five repeats of 10-fold cross validation, microbial variables were selected as the optimal predictor for CR (Supplementary Index 2) AUC 0.95 [95% CI 0.85-0.99] for FCCC, and AUC 0.88 [95%CI 0.77-0.94] for CW, and AUC 0.89 [95% CI 0.78-0.95] for combined data (Figure 4). Microbial markers for each institution depicted in supplementary index 2.

**Figure 4.**
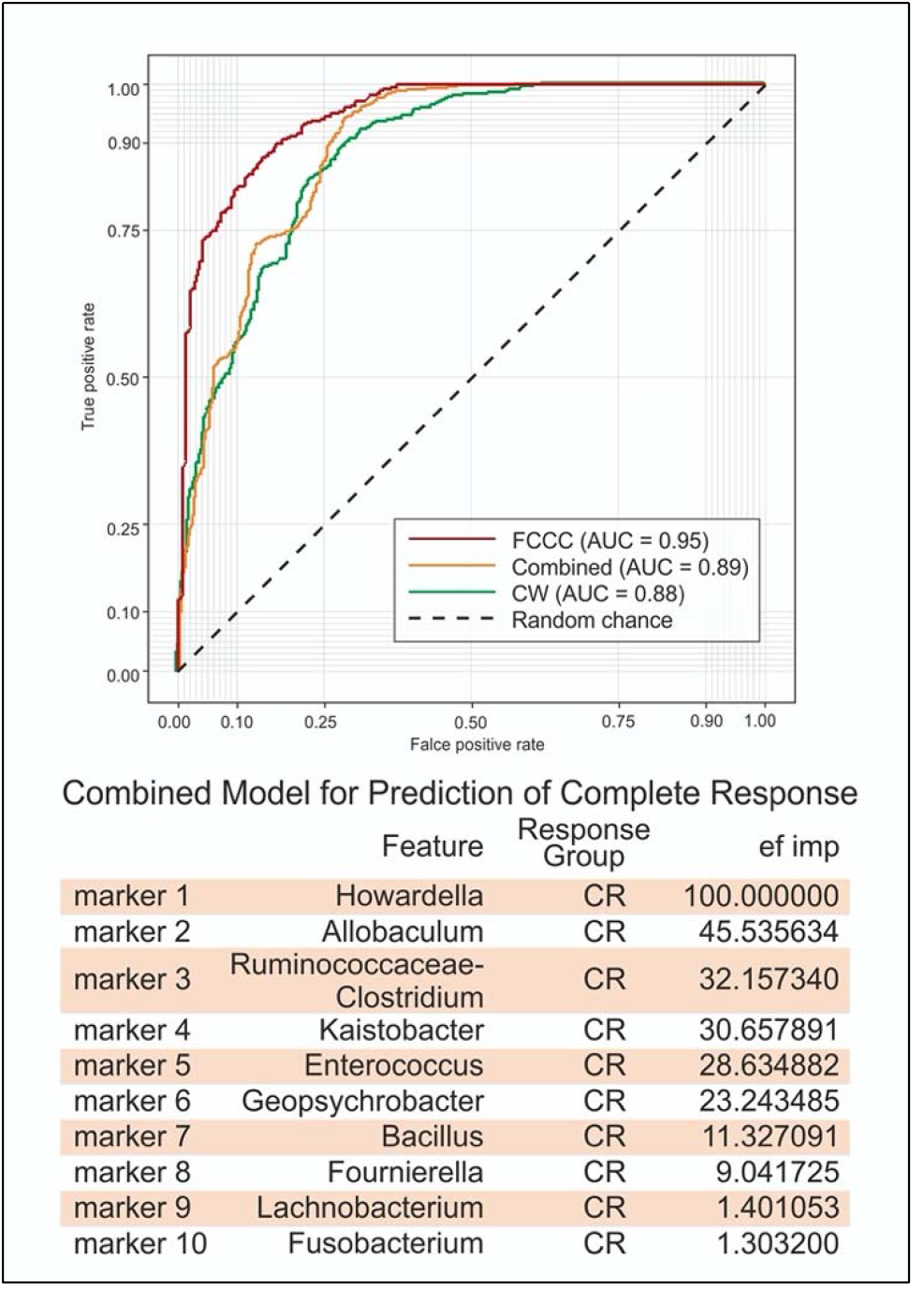
Prediction of NAC response status based on microbial biomarkers. The ROC curve for classifiers (OTUs) designated to discriminate between CR vs NR based on an LR model in CW, FCCC, and combined cohort. Individual markers (bacterial OTUs) for combined cohort and their feature importance score are listed in the table below AUC graph.

In addition to the microbial variables, a set of 24 clinical variables (Supplementary Index 2) were also utilized for variable selection to test for prediction for CR. Following identical LR modeling, we obtained an AUC 0.50 translating *to no discriminatory capability based on clinical and pathological factors alone for CR*.

### Microbiome composition during BBN-induced tumorigenesis

To measure GM changes associated with tumorigenesis, stool samples from mice exposed to BBN or regular drinking water were collected and compared at 1) baseline, 2) on treatment, 3) at end of BBN exposure, and 4) at the time that tumors were visualized radiographically followed by sacrifice to control for age-associated changes. We first analyzed the overall longitudinal changes within control and BBN-exposed mice, finding that there were no differences between control and BBN exposed mice at individual time points. (Figure 5 C, E, G). Similarly, alpha diversity differences between BBN across individual time points were not significant (pretreatment, 6, 12, 16-22 weeks). (Figure 5A) While beta diversity did show clustering based on time zero as compared to the remainder of time points (6,12,16-22 weeks), the differences were attributed to gut maturation rather than the exposure itself (Figure 5B).

**Figure 5.**
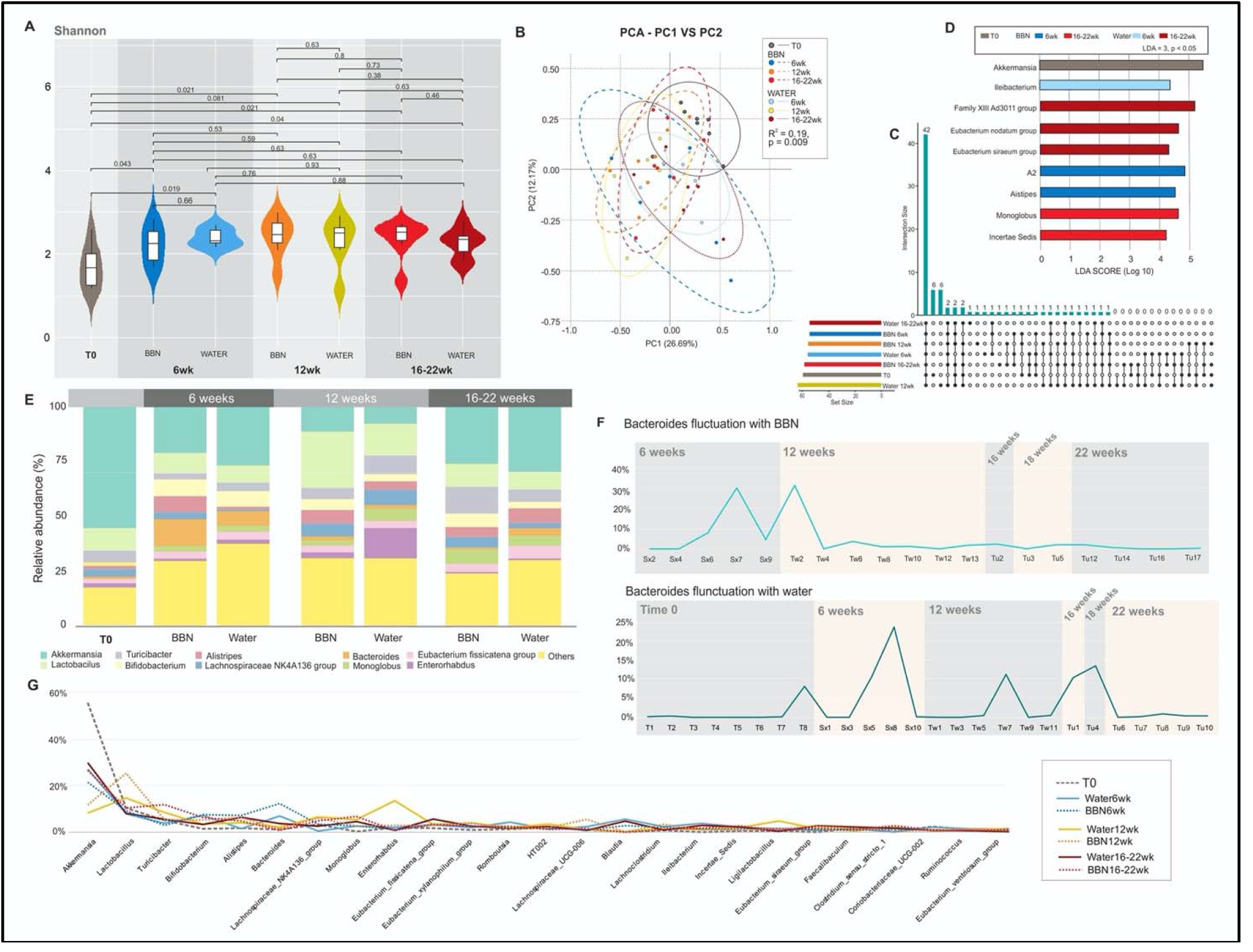
BBN vs water microbiota differences in murine model. (A) Alpha diversity index measurement by time (weeks) and treatment (BBN vs water), with beta diversity demonstrated in (B), showing no clustering, and low dissimilarity. (C) Upset plots depicting common and unique bacteria OTUs by treatment and week (D) LDA scores computed for differentially abundant taxa in GM of BBN vs water and time point of treatment, with length of bar indicating the effect size calculated. E. Bar graphs depicting phylogenic distribution of bacterial OTUs by treatment and time. (G) Direct comparison of bacterial OTUs by BBN vs water, and time of stool collection. (H) Fluctuation in Bacteroides prevalence depicted by genus and species over time

Due to the differences in *Bacteroides* noted in BC patients at both institutions, we examined *Bacteroides* abundance in both BBN exposed and water mice over the course of the experiment, noting overall increased prevalence of *Bacteroides* in BBN exposed mice (i.e 12.30% vs 6.89%, week 6) (Supplementary Index 1). Interestingly, at the end of BBN supplementation at 12 weeks, *Bacteroides* rapidly normalized to water GM prevalence (Figure 5F). LEfSe analysis highlighted an enrichment of *Monoglobus* in BBN exposed mice at 16-22 weeks, while water exposed mice were more likely to show enrichment of *Eubacterium*. Across both BBN and water exposed groups, higher *Akkermensia and Verrucomicrobiota* abundance was seen pretreatment (time zero) and declined thereafter at 6, 12, 16-22 weeks regardless of exposure (LDA 5, p<0.05; Supplementary Figure 5).

Due to sex dependent changes in microbial composition, we then analyzed compositional differences in microbiome in response to BBN exposure by sex.[18] No significant differences in alpha and beta diversity were seen in males or females separately across the timeline of the experiment (Supplementary Figure 6).

## Discussion

In this manuscript, we present the largest study to date describing an association between the GM and NAC response in MIBC. In addition to characterizing the differences between CTR and BC patients, we extended our research to analyze fluctuations of gut microbiome throughout NAC. Furthermore, to assess for potential environmental confounding, we analyzed the changes of fecal microbiome of the patients’ partners throughout treatment. Using microbial data from both institutions, we were able to construct a classifier for CR prediction, and test its accuracy compared to clinicopathologic features alone. Moreover, we assessed gut microbiome changes with BBN exposure in our murine model to characterize the variability of the microbiome during tumorigenesis. As bacterial composition influences key metabolic and immunologic functions in health and disease, understanding time point and microbiome specific alterations during the tumorigenesis, can provide additional data for surveillance of disease and recurrence.

Our results are consistent with previous findings of associations of GM in tumorigenesis. In particular, our cohorts highlighted an increase in *Prevotella, Fusobacteria*, and *Bacteroides* in BC patients, all widely reported to be associated both with tumorigenesis and response to therapy [19-22]. In addition to individual associations, our real-world cohorts from both institutions showed similar clustering patterns across communities (groups I-III) and association with chemotherapy response. More specifically, both institutions displayed communities that were *Bacteroides*-dominant or *Prevotella* dominant. <u>Although these communities are not distinct and represent a continuous gradient, our results indicate strong association of residual disease in patients with</u> *<u>Bacteroides</u>*<u>-dominant community across both institutions, regardless of NAC regimen and geography.</u>

There has been a rapid development of novel cancer therapies, however, multiple challenges in understanding the host-tumor interaction and prediction of treatment success remain. One is unable to fully elucidate the inter-individual response differences based on somatic genetic variants, or clinical features alone, implying presence of other/undescribed factors contributing therapeutic susceptibility. The concept of the GM as the “pharmacomicrobiomics” focuses specifically on the interplay of drug modification, distribution, and biological pathways that impact therapeutic outcome may be relevant to cancer treatment. A growing body of literature has revealed the intimate relationship between the gut microbiome and therapeutic response[23, 24], harnessing the microbiota to optimize cancer treatment has become an alternative avenue for personalized medicine.

As bacterial metabolism of the drug can significantly reduce the chemotherapeutic efficacy via enzymatic reaction, it is not surprising that our cohort of patients treated with gemcitabine-cisplatin and higher abundance of E.coli were more likely to show residual disease. While intrinsic gemcitabine chemoresistance of the tumor may be present, recently published evidence points to enzymatic detoxification of gemcitabine by cytidine deaminase enzyme present in gammaproteobacteria such as E.coli.[25] It is uncertain what microbial biomass is required to inactive gemcitabine, or if gammaproteobacteria in the gut could detoxify the drug, but our findings are consistent with this effect. We did not measure gemcitabine deamination byproducts within stool or blood, however, early preclinical studies are underway translating these mechanistic findings into clinical applications.

Our study clearly shows distinct GM composition is associated with chemotherapy response. More specifically, the microbes enriched in responders (*Howardella and Prevotella)* are both butyrate producers, a type of short chain fatty acid (SCFA). This SCFA promotes regulatory T cell function and anti-inflammatory response by binding to G-protein coupled receptor 109a expressed on myeloid cells and colonocytes [26]. SFCA, specifically butyrate, are also histone deacetylase inhibitors and can positively redirect expression programs within tumors. [27] The key finding from our analysis is the association of *Bacteroides* with poor NAC response in both GC and ddMVAC cohorts. The relationship between *Bacteroides* and poor treatment is not unique to bladder cancer; minimal benefit of both immunotherapy and chemotherapy has been observed in melanoma [9], pancreatic [28], and colon cancers [26].

The dynamic variation of the GM over the course of NAC in bladder cancer has not been studied, however, studies within colorectal[29], head and neck[30] and cervical cancer [31] [32]have all demonstrated significant fluctuations in composition. It is therefore not surprising, that the longitudinal microbiota fluctuations in our study are mostly driven by an increase in *Bacteroides abundance* during the course of chemotherapy. Unique to our cohort is the inclusion of patient’s partners as means to control for environmental confounders. Unlike the patients who exhibited a rise in *Bacteroides* at the end of treatment, their respective live-in partners did not show this synchronous shift. These results help affirm the findings of a true microbiota shift, rather than spurious fluctuation.

It is important to also highlight an interesting yet still controversial notion of existence of “enterotypes” characterized by dominant genera (*Bacteroides, Prevotella and Ruminococcus)*, and their co-occurring dietary traits, first published by Arumugam et al [33]. In particular, the *Bacteroides* enterotype was found to be associated with high animal protein and saturated fat, while the *Prevotella* enterotype was observed with high fiber/plant-based nutrition (low meat). As there is growing awareness of importance of dietary phenotype with health and disease and recognition of host-microbiome mutualism, it is difficult to say whether the dietary phenotype in our study (high fiber/plant based *Prevotella* subtype) and response to chemotherapy is result of potentially drug targetable intervention or confounding by an environmental factor.

Although animal models are widely used to investigate human disease on molecular and genetic level, murine microbiota models are relatively complex due to the effects of the environment. Nonetheless, gut microbiome of mice and humans are similar in that they are both made up of roughly 90% Firmicutes and Bacteroidetes [34]. Although this similarity may seem indicative of high gut microbiota resemblance, the most discernible difference is the ratio of the two major phyla, with humans having a greater Firmicutes/Bacteroidetes ratio, whereas the inverse is true for mice [34, 35]. Such differences in the Firmicutes/Bacteriodetes ratio does imply a limitation to the use of these animal models for *association testing* since the populations of major groups belonging to these phyla might not imitate the ratio typically found in a healthy human gut. Our study, however, aimed to explore longitudinal changes of mice exposed to a carcinogen, hoping to identify a key timepoint of transformation and dysbiosis. Mice exposed to BBN in our study similarly showed a steady increase in *Bacteroides* up to week 12 when BBN was stopped, with a precipitous drop following withdrawal of the carcinogen. The mice never received treatment for their BBN induced tumors, thus we cannot extrapolate the implication of this finding. It does, however, provide evidence regarding the feasibility of *Bacteroides* manipulation in murine model, particularly in future models of therapeutics.

While it is well known that *Bacteroides* and *E. coli* possess a unique ability for pathogenic invasion and biofilm formation generating carcinogenic metabolic byproducts such as sialic acid, fucose, and succinate [36]the mechanism by which *Bacteroides affects* chemotherapy response remains unclear. It is evident, however, that inhibition of *Bacteroides* during treatment with microbiome modulation such as *Lactobacillus* administration (*Bacteroides* inhibitor), antibiotics aimed to target *Bacteroides*, or whole fecal microbiota transplantation represent an area of significant research interest, and applicability to many solid tumors. [37]

Although in line with previous research in other solid tumors, enrichment of certain microbes in responders or non-responders in our study does not indicate a direct association, rather a correlation. The microbial signature, however, outperforms a set of clinical variables (without tumor genetic variables) for prediction of CR (AUC 0.89). It is clear that our classifier performs robustly in screening out responders, which may facilitate personalized treatment decisions.

Our results suggest that baseline gut microbiota as well as evolving fluctuations throughout chemotherapy are important determinants of CR. Further studies are needed to could synthesize analysis of the microbial signature, clinical variables, genomic classifiers, and circulating tumor DNA to develop stronger predictive/prognostic biomarkers.

One of the main objectives of our research is to identify patients who will not benefit from NAC but who may benefit from an alternative treatment approach or early surgery. Furthermore, by beginning to describe a predictive microbial signature, larger studies to validate the findings or preclinical studies to evaluate the effect of GM modulation on chemoresponse could one day support clinical trials of GM modulation for therapeutic benefit as has already been undertaken in melanoma patients

Our study is not without limitations, including small sample size, and variable treatments between the institutions. Future studies could also assess the peripheral and intratumor immune landscape as it relates to favorable or unfavorable GM. Another limitation is lack of concurrent tumor data including genomic mutations, and immune infiltration for response classifier. However, this study is unique as it analyzes separate cohorts, with similar findings independent from geography and chemotherapy regimen. Furthermore, inclusion of partners of our patients allowed to adjust for environmental exposure that may alter microbiome composition.

In conclusion, we found that a set of microbial features in the baseline GM of bladder cancer patients is associated with response to chemotherapy. This is supported by both cohorts, highlighting higher abundance of *Bacteroides* as a poor prognostic factor for CR. The data presented above warrants further validation in large prospective trial, combined with immune and genomic markers of the host and tumor. We hope these preliminary results are encouraging and lay the groundwork for further ongoing trials

## Supporting information

Supplemental Index 1

Supplemental Index 2

Supplemental Index 3

## Abbreviations

NGS: Next-Generation Sequencing
BC: Bladder Cancer
CTR: Controls
rRNA: Ribosomal RNA
OTU: Operational Taxonomic Unit
QIIME: Quantitative Insights into Microbial Ecology
PCoA: Principal Coordinate Analysis
LEfSe: Linear discriminant analysis Effect Size
CLR: Centered Log Ratios
NR: Non-Responder
CR: Complete Response
CTRL: Controls
MIBC: Muscle-Invasive Bladder Cancer
NMIBC: Non-Muscle Invasive Bladder Cancer
FDR: False Discovery Rate
ctDNA: Circulating tumor DNA
NAC: Neoadjuvant chemotherapy
GM: Gut microbiome

## Supplementary Figure

**Supplementary Figure 1:**
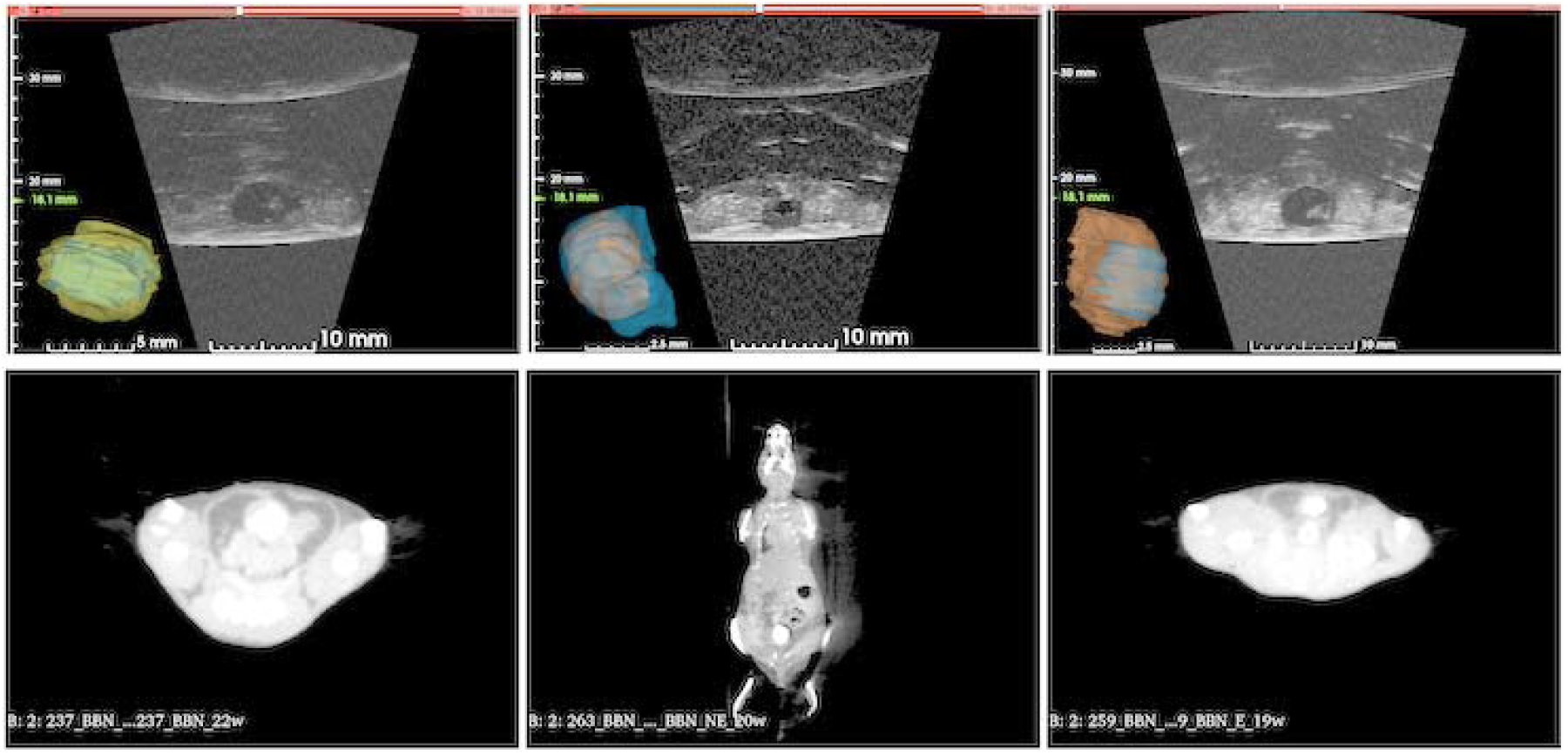
CT urography and US imaging of mice showing BBN induced urothelial cancer in mouse in axial and coronal views

**Supplementary Figure 2:**
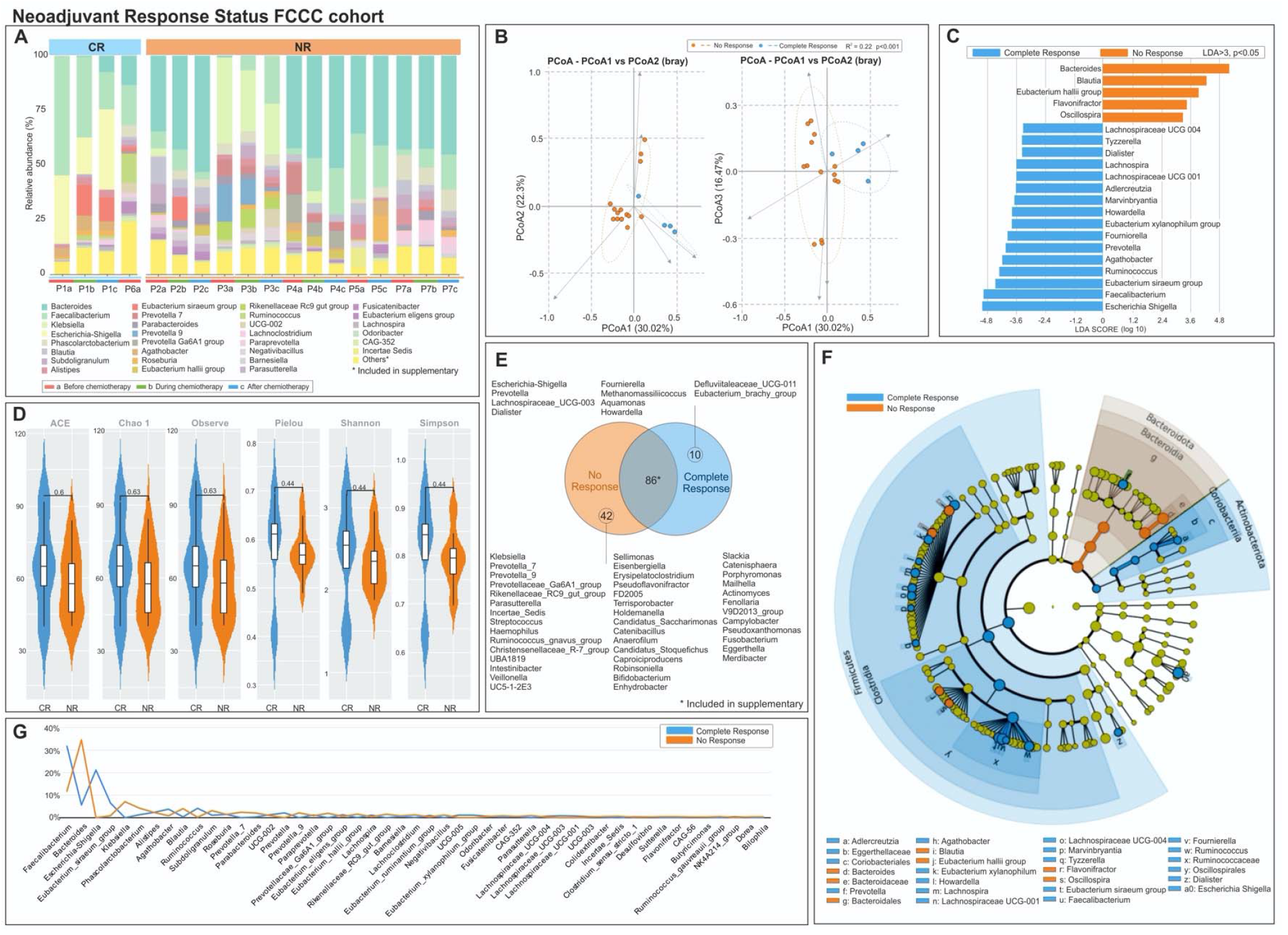
Differences in gut microbiome by pathological response to neoadjuvant chemotherapy. A) Stacked bar plots of individual patients designated as letter and subsequent collection timing (1: before chemo, 2: during chemo, 3: after chemo) and composition of common bacteria by response status. (B) Clustering of responders vs non responder based on beta diversity of GM in stool sample at time of collection (C)LDA scores for differentially abundant taxa in CR vs NR, length of bar indicating effect size. (D) Alpha diversity indices between responder status (E) Venn diagram depicting number of shared and unique bacterial OTUs based on NAC response status with additional information included in supplementary index 3. (F) Taxonomic cladogram depicting differences in GM from LEfSe (G) Overall abundance distribution differences in CR vs NR.

**Supplementary Figure 3:**
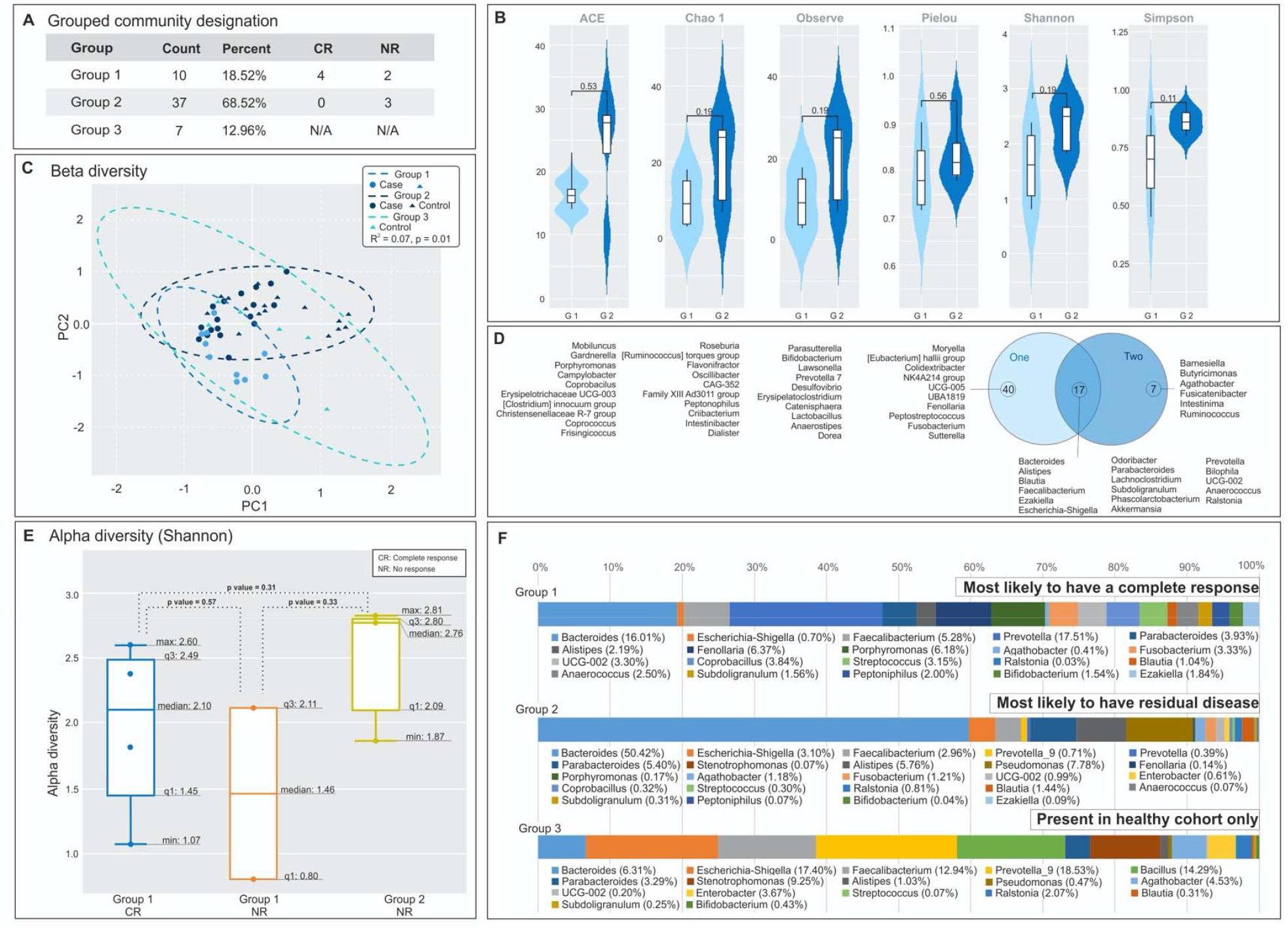
Classification of response status based on community grouping at CW: A. Identification of three groups based on hierarchal clustering, and distribution of patients (CR and NR) vs controls. Additional information included in supplementary index 1. (B) Alpha and (C) Beta diversity showing significant differences in bacterial OTUs and clustering among the community groups. (D) Further unique and shared bacterial OTUs in group I and II (groups with BC patients) and their respective differences in alpha diversity (E). Furthermore, as depicted in (E), there were zero patients with CR in Group II. (F) Group specific abundances are depicted as phylogenetic columns.

**Supplementary Figure 4:**
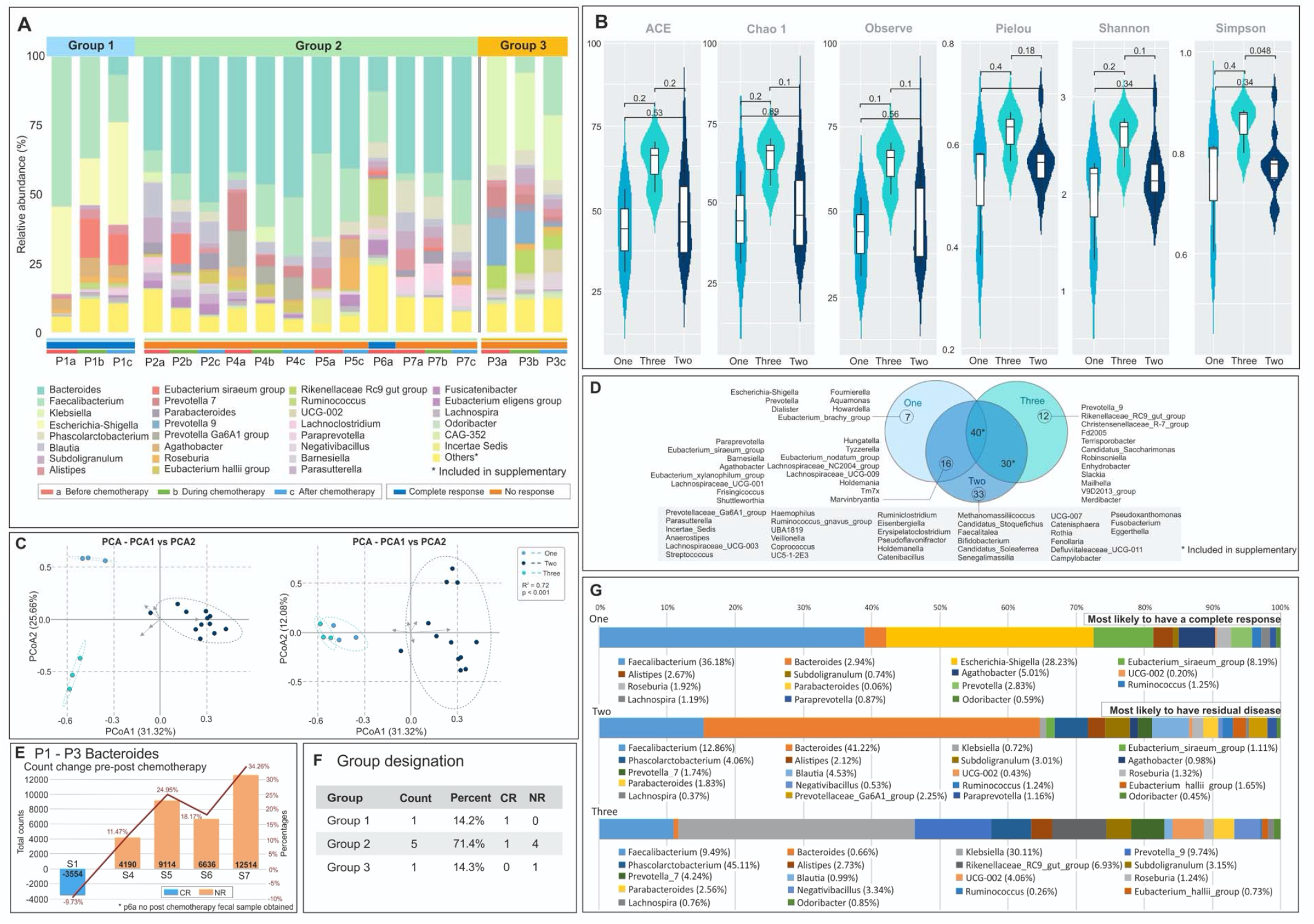
Classification of response status based on community grouping at FCCC. A. Identification of three groups based on hierarchal clustering, and distribution of patients (CR and NR) vs controls. Additional information included in supplementary index 1. (B) Alpha and (C) Beta diversity showing significant differences in bacterial OTUs and clustering among the community groups. (D) Further unique and shared bacterial OTUs in group I and II (groups with BC patients) and their respective differences in alpha diversity (E). Furthermore, as depicted in (E), there were zero patients with CR in Group II. (F) Group specific abundances are depicted as phylogenetic columns.

**Supplementary Figure 5:**
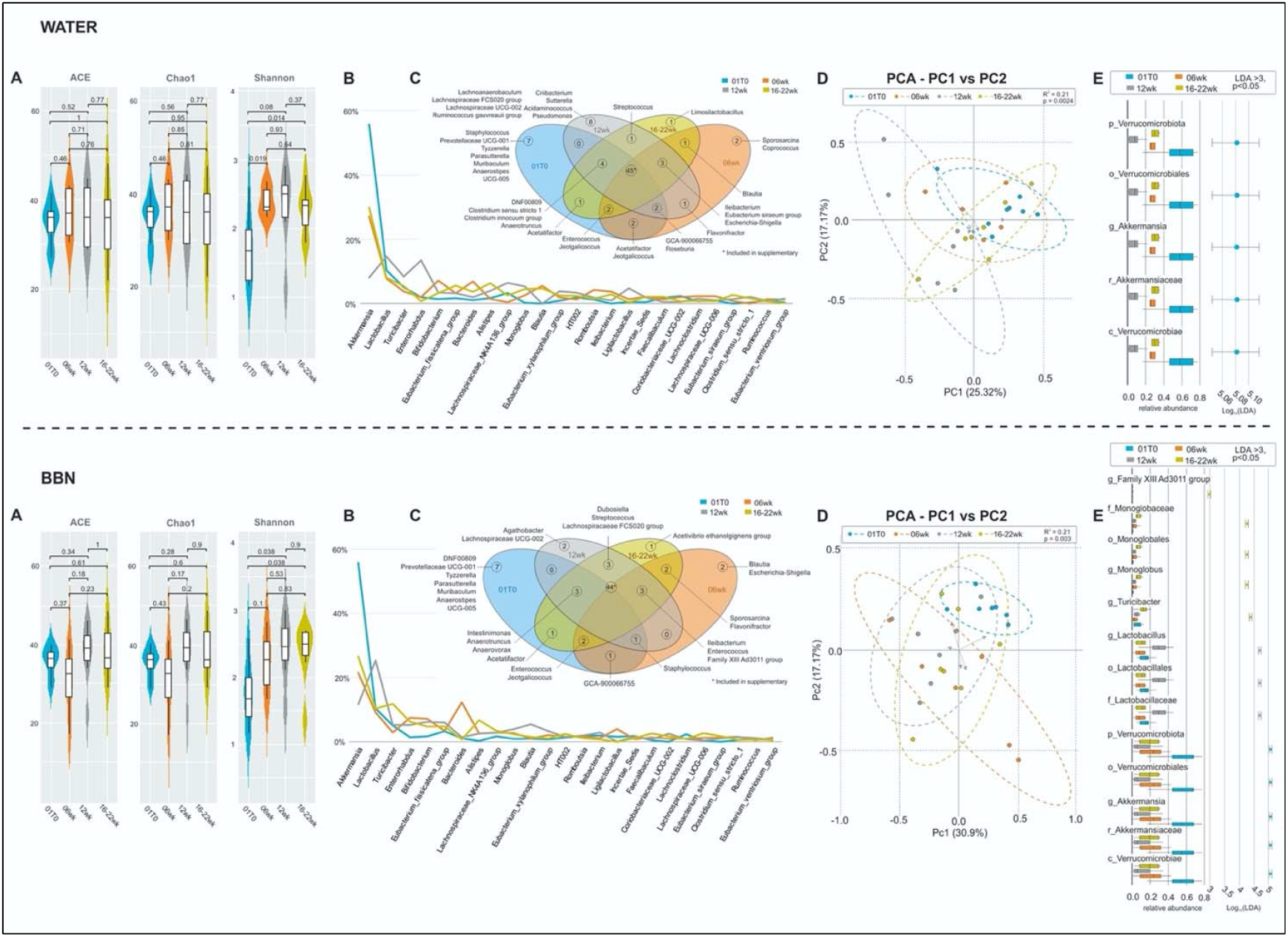
Differences in microbial composition of the gut in BBN vs water in mice by weeks. (A) Alpha diversity indices differences in mice exposed to BBN (top) or Water (bottom) at baseline (0 weeks) compared to 6, 12, 18-22 weeks showing no significant difference. (B) Microbial abundance by individual time points and (C) Venn diagram depiction of unique and common bacterial OTUs from baseline to week 22. (D) Beta diversity, although dissimilar to baseline, no otherwise significant clustering was seen by week 22. (E) LDA scores computed for differentially abundant taxa by time point, demonstrating individual taxa differences by taxonomic classification up to genus.

**Supplementary Figure 6:**
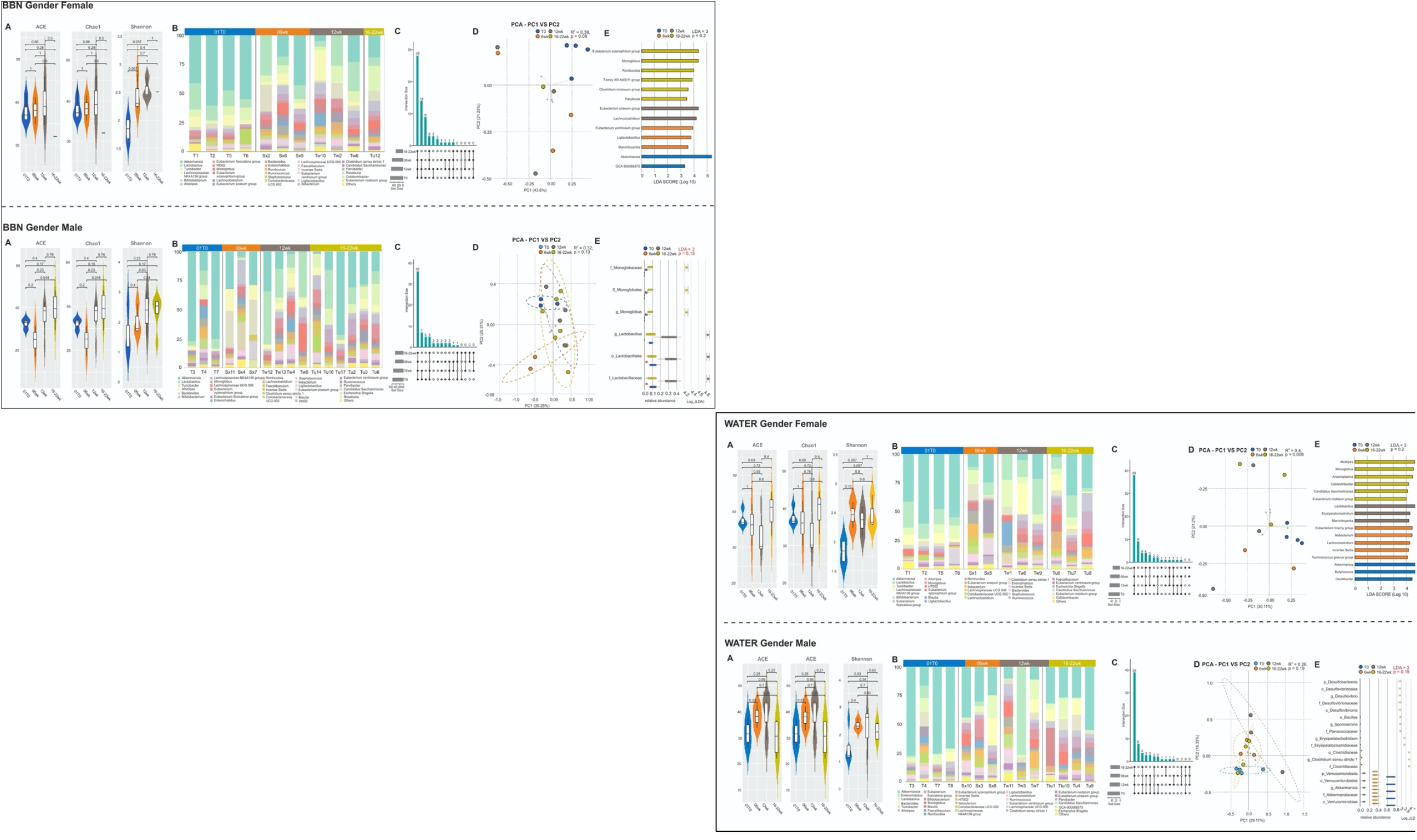
Differences in microbial composition of the gut in BBN vs water in mice by sex. (A) Alpha diversity indices differences in mice exposed to BBN (top) or Water (bottom) at baseline (0 weeks) compared to 6, 12, 18-22 weeks by sex showing no detectable difference (B) Microbial abundance by individual time points and (C) Upset plots depiction of unique and common bacterial OTUs from baseline to week 22 by sex (D) Beta diversity with no minimal dissimilarity among group types regardless of sex (E) LDA scores computed for differentially abundant taxa by time point, demonstrating individual taxa differences by taxonomic classification up to genus.

## Notes

**Source of Funding:** P30CA043703 Case Western Comprehensive Cancer Center Microbiome Grant, P30 CA006927 Fox Chase Cancer Center Support Grant, NIH grant # R01AI145289-01A1 (MG), CA181178 Department of Defense CDMRP (PHA).

### Competing Interest Statement

The authors have declared no competing interest.

## References

1. Rouprêt, M., et al., Oncologic outcomes and survival in pT0 tumors after radical cystectomy in patients without neoadjuvant chemotherapy: results from a large multicentre collaborative study. Ann Surg Oncol, 2011. 18(13): p. 3833–8.

2. Petrelli, F., et al., Correlation of pathologic complete response with survival after neoadjuvant chemotherapy in bladder cancer treated with cystectomy: a meta-analysis. Eur Urol, 2014. 65(2): p. 350–7.

3. Plimack, E.R., et al., Defects in DNA Repair Genes Predict Response to Neoadjuvant Cisplatin-based Chemotherapy in Muscle-invasive Bladder Cancer. Eur Urol, 2015. 68(6): p. 959–67.

4. Liu, D., et al., Clinical Validation of Chemotherapy Response Biomarker ERCC2 in Muscle-Invasive Urothelial Bladder Carcinoma. JAMA Oncol, 2016. 2(8): p. 1094–6.

5. Lee, K.A., et al., The gut microbiome: what the oncologist ought to know. Br J Cancer, 2021. 125(9): p. 1197–1209.

6. Huang, J., et al., Effects of microbiota on anticancer drugs: Current knowledge and potential applications. EBioMedicine, 2022. 83: p. 104197.

7. Consortium, H.M.P., A framework for human microbiome research. Nature, 2012. 486(7402): p. 215–21.

8. Lundberg, D.S., et al., Practical innovations for high-throughput amplicon sequencing. Nat Methods, 2013. 10(10): p. 999–1002.

9. Gopalakrishnan, V., et al., Gut microbiome modulates response to anti-PD-1 immunotherapy in melanoma patients. Science, 2018. 359(6371): p. 97–103.

10. Yang, T.W., et al., Enterotype-based Analysis of Gut Microbiota along the Conventional Adenoma-Carcinoma Colorectal Cancer Pathway. Sci Rep, 2019. 9(1): p. 10923.

11. Yilmaz, P., et al., The SILVA and “All-species Living Tree Project (LTP)” taxonomic frameworks. Nucleic Acids Res, 2014. 42(Database issue): p. D643–8.

12. Bolyen, E., et al., Reproducible, interactive, scalable and extensible microbiome data science using QIIME 2. Nat Biotechnol, 2019. 37(8): p. 852–857.

13. Segata, N., et al., Metagenomic biomarker discovery and explanation. Genome Biol, 2011. 12(6): p. R60.

14. Costea, P.I., et al., Enterotypes in the landscape of gut microbial community composition. Nat Microbiol, 2018. 3(1): p. 8–16.

15. computing, R.C.T.R.A.l.a.e.f.s. R foundation for Statistical Computing: Austria.

16. (2015), P.C.T., Python: A dynamic, open source programming language.

17. Cao, Y., et al., microbiomeMarker: an R/Bioconductor package for microbiome marker identification and visualization. Bioinformatics, 2022. 38(16): p. 4027–4029.

18. Elderman, M., et al., Sex and strain dependent differences in mucosal immunology and microbiota composition in mice. Biol Sex Differ, 2018. 9(1): p. 26.

19. Shariati, A., et al., Association between colorectal cancer and Fusobacterium nucleatum and Bacteroides fragilis bacteria in Iranian patients: a preliminary study. Infect Agent Cancer, 2021. 16(1): p. 41.

20. Png, C.W., et al., Alterations in co-abundant bacteriome in colorectal cancer and its persistence after surgery: a pilot study. Sci Rep, 2022. 12(1): p. 9829.

21. Sevcikova, A., et al., The Impact of the Microbiome on Resistance to Cancer Treatment with Chemotherapeutic Agents and Immunotherapy. Int J Mol Sci, 2022. 23(1).

22. Serna, G., et al., Fusobacterium nucleatum persistence and risk of recurrence after preoperative treatment in locally advanced rectal cancer. Ann Oncol, 2020. 31(10): p. 1366–1375.

23. Chrysostomou, D., et al., Gut Microbiota Modulation of Efficacy and Toxicity of Cancer Chemotherapy and Immunotherapy. Gastroenterology, 2022.

24. Ting, N.L., H.C. Lau, and J. Yu, Cancer pharmacomicrobiomics: targeting microbiota to optimise cancer therapy outcomes. Gut, 2022. 71(7): p. 1412–1425.

25. Geller, L.T., et al., Potential role of intratumor bacteria in mediating tumor resistance to the chemotherapeutic drug gemcitabine. Science, 2017. 357(6356): p. 1156–1160.

26. Chaput, N., et al., Baseline gut microbiota predicts clinical response and colitis in metastatic melanoma patients treated with ipilimumab. Ann Oncol, 2019. 30(12): p. 2012.

27. Fellows, R., et al., Microbiota derived short chain fatty acids promote histone crotonylation in the colon through histone deacetylases. Nat Commun, 2018. 9(1): p. 105.

28. Del Castillo, E., et al., The Microbiomes of Pancreatic and Duodenum Tissue Overlap and Are Highly Subject Specific but Differ between Pancreatic Cancer and Noncancer Subjects. Cancer Epidemiol Biomarkers Prev, 2019. 28(2): p. 370–383.

29. Alexander, J.L., et al., Gut microbiota modulation of chemotherapy efficacy and toxicity. Nat Rev Gastroenterol Hepatol, 2017. 14(6): p. 356–365.

30. Bahig, H., et al., Longitudinal characterization of the tumoral microbiome during radiotherapy in HPV-associated oropharynx cancer. Clin Transl Radiat Oncol, 2021. 26: p. 98–103.

31. El Alam, M.B., et al., A prospective study of the adaptive changes in the gut microbiome during standard-of-care chemoradiotherapy for gynecologic cancers. PLoS One, 2021. 16(3): p. e0247905.

32. Sims, T.T., et al., Gut microbiome diversity is an independent predictor of survival in cervical cancer patients receiving chemoradiation. Commun Biol, 2021. 4(1): p. 237.

33. Arumugam, M., et al., Enterotypes of the human gut microbiome. Nature, 2011. 473(7346): p. 174–80.

34. Krych, L., et al., Quantitatively different, yet qualitatively alike: a meta-analysis of the mouse core gut microbiome with a view towards the human gut microbiome. PLoS One, 2013. 8(5): p. e62578.

35. Nagpal, R., et al., Comparative Microbiome Signatures and Short-Chain Fatty Acids in Mouse, Rat, Non-human Primate, and Human Feces. Front Microbiol, 2018. 9: p. 2897.

36. Ivleva, E.A. and S.I. Grivennikov, Microbiota-driven mechanisms at different stages of cancer development. Neoplasia, 2022. 32: p. 100829.

37. Wexler, H.M., Bacteroides: the good, the bad, and the nitty-gritty. Clin Microbiol Rev, 2007. 20(4): p. 593–621.

