## Supplemental Index 3 for "Role of Gut Microbiome in Neoadjuvant Chemotherapy Response in Urothelial Carcinoma: A Multi-Institutional Prospective Cohort Evaluation"

CW Group Community Venn Diagram

| |
|---|
| **N=40** |
| Faecalibacterium |
| UCG-002 |
| Subdoligranulum |
| Alistipes |
| Parabacteroides |
| CAG-352 |
| Roseburia |
| Desulfovibrio |
| Dorea |
| Odoribacter |
| Clostridium_sensu_stricto_1 |
| UCG-003 |
| CAG-56 |
| Lachnospira |
| Colidextribacter |
| Bacteroides |
| Sutterella |
| Lachnospiraceae_UCG-004 |
| Butyricimonas |
| Ruminococcus |
| Romboutsia |
| Flavonifractor |
| Monoglobus |
| Lachnoclostridium |
| Paludicola |
| Butyricicoccus |
| Ruminococcus_gauvreauii_group |

| |
|---|
| Bilophila |
| Peptococcus |
| Lachnospiraceae_ND3007_group |
| Oxalobacter |
| Intestinimonas |
| DTU089 |
| Anaerotruncus |
| Adlercreutzia |
| Erysipelotrichaceae_UCG-003 |
| Eubacterium_eligens_group |
| Harryflintia |
| Angelakisella |
| Lachnospiraceae_UCG-010 |
| **N=30** |
| Klebsiella |
| Phascolarctobacterium |
| Prevotella_7 |
| Negativibacillus |
| Eubacterium_ruminantium_group |
| Blautia |
| Eubacterium_hallii_group |
| Intestinibacter |
| Oscillospira |
| Fusicatenibacter |
| Ruminococcus_torques_group |
| Family_XIII_AD3011_group |
| Coprobacter |

| |
|---|
| UCG-005 |
| Oscillibacter |
| Hespellia |
| NK4A214_group |
| GCA-900066575 |
| Sanguibacteroides |
| Tuzzerella |
| Phocea |
| Lachnospiraceae_FCS020_group |
| Collinsella |
| Anaerofilum |
| Caproiciproducens |
| UCG-009 |
| Porphyromonas |
| Sellimonas |
| Cloacibacillus |
| Actinomyces |

FCCC cohort Venn Diagram during chemotherapy

| |
|---|
| N=102 |
| [Eubacterium]_eligens_group |
| [Eubacterium]_hallii_group |
| [Eubacterium]_nodatum_group |
| [Eubacterium]_ruminantium_group |
| [Eubacterium]_siraeum_group |
| [Eubacterium]_xylanophilum_group |
| [Ruminococcus]_gauvreauii_group |
| [Ruminococcus]_gnavus_group |
| [Ruminococcus]_torques_group |
| Adlercreutzia |
| Agathobacter |
| Alistipes |
| Anaerofilum |
| Anaerostipes |
| Anaerotruncus |
| Angelakisella |
| Bacteroides |
| Barnesiella |
| Bilophila |
| Blautia |
| Butyricicoccus |
| Butyricimonas |
| CAG-352 |
| CAG-56 |
| Candidatus_Soleaferrea |
| Candidatus_Stoquefichus |
| Caproiciproducens |

| |
|---|
| Catenibacillus |
| Cloacibacillus |
| Clostridium_sensu_stricto_1 |
| Colidextribacter |
| Collinsella |
| Coprobacter |
| Desulfovibrio |
| Dorea |
| DTU089 |
| Eisenbergiella |
| Erysipelatoclostridium |
| Erysipelotrichaceae_UCG-003 |
| Escherichia-Shigella |
| Faecalibacterium |
| Faecalitalea |
| Family_XIII_AD3011_group |
| FD2005 |
| Flavonifractor |
| Frisingicoccus |
| Fusicatenibacter |
| GCA-900066575 |
| Harryflintia |
| Hespellia |
| Holdemania |
| Howardella |
| Hungatella |
| Intestinibacter |
| Intestinimonas |

| |
|---|
| Klebsiella |
| Lachnoclostridium |
| Lachnospira |
| Lachnospiraceae_FCS020_group |
| Lachnospiraceae_NC2004_group |
| Lachnospiraceae_ND3007_group |
| Lachnospiraceae_UCG-001 |
| Lachnospiraceae_UCG-004 |
| Lachnospiraceae_UCG-010 |
| Marvinbryantia |
| Monoglobus |
| Negativibacillus |
| Odoribacter |
| Oscillibacter |
| Oscillospira |
| Oxalobacter |
| Paludicola |
| Parabacteroides |
| Paraprevotella |
| Parasutterella |
| Peptococcus |
| Phascolarctobacterium |
| Phocea |
| Prevotella |
| Prevotella_7 |
| Prevotella_9 |
| Prevotellaceae_Ga6A1_group |
| Pseudoflavonifractor |

| |
|---|
| Rikenellaceae_RC9_gut_group |
| Robinsoniella |
| Romboutsia |
| Roseburia |
| Ruminiclostridium |
| Ruminococcus |
| Sanguibacteroides |
| Sellimonas |
| Senegalimassilia |
| Shuttleworthia |
| Subdoligranulum |
| Sutterella |
| TM7x |
| Tyzzerella |
| UBA1819 |
| UC5-1-2E3 |
| UCG-002 |
| UCG-003 |
| Veillonella |

FCCC Group Community Venn Diagram

| |
|---|
| **N=40** |
| Faecalibacterium |
| UCG-002 |
| Subdoligranulum |
| Alistipes |
| Parabacteroides |
| CAG-352 |
| Roseburia |
| Desulfovibrio |
| Dorea |
| Odoribacter |
| Clostridium_sensu_stricto_1 |
| UCG-003 |
| CAG-56 |
| Lachnospira |
| Colidextribacter |
| Bacteroides |
| Sutterella |
| Lachnospiraceae_UCG-004 |
| Butyricimonas |
| Ruminococcus |
| Romboutsia |
| Flavonifractor |
| Monoglobus |
| Lachnoclostridium |
| Paludicola |
| Butyricicoccus |
| Ruminococcus_gauvreauii_group |

| |
|---|
| Bilophila |
| Peptococcus |
| Lachnospiraceae_ND3007_group |
| Oxalobacter |
| Intestinimonas |
| DTU089 |
| Anaerotruncus |
| Adlercreutzia |
| Erysipelotrichaceae_UCG-003 |
| Eubacterium_eligens_group |
| Harryflintia |
| Angelakisella |
| Lachnospiraceae_UCG-010 |
| **N=30** |
| Klebsiella |
| Phascolarctobacterium |
| Prevotella_7 |
| Negativibacillus |
| Eubacterium_ruminantium_group |
| Blautia |
| Eubacterium_hallii_group |
| Intestinibacter |
| Oscillospira |
| Fusicatenibacter |
| Ruminococcus_torques_group |
| Family_XIII_AD3011_group |
| Coprobacter |

| |
|---|
| UCG-005 |
| Oscillibacter |
| Hespellia |
| NK4A214_group |
| GCA-900066575 |
| Sanguibacteroides |
| Tuzzerella |
| Phocea |
| Lachnospiraceae_FCS020_group |
| Collinsella |
| Anaerofilum |
| Caproiciproducens |
| UCG-009 |
| Porphyromonas |
| Sellimonas |
| Cloacibacillus |
| Actinomyces |

| | |
|---|---|
| P1a | |
| Prevotella | 2.54% |
| Clostridium_sensu_stricto_1 | 1.28% |
| P1b | |
| Prevotella | 5.03% |
| Eubacterium_xylanophilum_group | 2.19% |
| Lachnospiraceae_UCG-001 | 1.13% |
| P1c | |
| UCG-003 | 1.37% |
| Lachnospiraceae_UCG-001 | 1.22% |
| Eubacterium_xylanophilum_group | 1.20% |
| P2a | |
| Haemophilus | 2.26% |
| Streptococcus | 2.00% |
| Colidextribacter | 1.55% |
| Ruminococcus_torques_group | 1.47% |
| Bilophila | 1.06% |
| P2b | |
| Bilophila | 2.22% |
| Butyricimonas | 1.10% |
| P2c | |
| - | |
| P3a | |
| Desulfovibrio | 1.99% |
| Dorea | 1.27% |
| P3b | |
| UCG-003 | 1.60% |
| Dorea | 1.25% |
| Clostridium_sensu_stricto_1 | 1.25% |
| Desulfovibrio | 1.15% |
| Colidextribacter | 1.13% |
| P3c | |
| Eubacterium_ruminantium_group | 3.87% |
| Christensenellaceae_R-7_group | 1.07% |
| P4a | |
| Lachnospiraceae_UCG-004 | 1.77% |
| Anaerostipes | 1.77% |
| Ruminococcus_gauvreauii_group | 1.54% |
| Romboutsia | 1.31% |
| P4b | |
| Ruminococcus_gauvreauii_group | 2.36% |
| Tuzzerella | 1.97% |
| Ruminococcus_torques_group | 1.09% |
| P4c | |
| - | |
| P5a | |
| - | |
| P5c | |
| Romboutsia | 1.25% |
| P6a | |
| UCG-005 | 4.59% |
| Eubacterium_ruminantium_group | 3.92% |
| Lachnospiraceae_UCG-003 | 3.45% |
| NK4A214_group | 1.88% |
| Desulfovibrio | 1.25% |
| Lachnospiraceae_UCG-004 | 1.16% |
| Eubacterium_xylanophilum_group | 1.04% |
| P7a | |
| Sutterella | 2.03% |
| Oscillibacter | 1.33% |
| Flavonifractor | 1.31% |
| Coprobacter | 1.25% |
| P7b | |
| Oscillibacter | 1.80% |
| Sutterella | 1.46% |
| P7c | |
| Sutterella | 1.57% |

FCC NAC response Venn Diagram

| |
|---|
| N=86 |
| Faecalibacterium |
| Bacteroides |
| Phascolarctobacterium |
| Blautia |
| Subdoligranulum |
| Alistipes |
| Parabacteroides |
| Eubacterium_hallii_group |
| Roseburia |
| Lachnoclostridium |
| Negativibacillus |
| UCG-002 |
| Eubacterium_siraeum_group |
| Paraprevotella |
| Agathobacter |
| Fusicatenibacter |
| Barnesiella |
| CAG-352 |
| Odoribacter |
| Eubacterium_eligens_group |
| Colidextribacter |
| Ruminococcus_gauvreauii_group |
| Flavonifractor |
| Sutterella |
| Lachnospira |
| Oscillibacter |
| Bilophila |

| |
|---|
| Dorea |
| Oscillospira |
| CAG-56 |
| Ruminococcus_torques_group |
| Eubacterium_ruminantium_group |
| Butyricimonas |
| Desulfovibrio |
| Lachnospiraceae_UCG-004 |
| Anaerostipes |
| Romboutsia |
| UCG-003 |
| Coprobacter |
| Clostridium_sensu_stricto_1 |
| Ruminococcus |
| Butyricicoccus |
| Tuzzerella |
| Intestinimonas |
| Monoglobus |
| Anaerotruncus |
| Lachnospiraceae_UCG-010 |
| GCA-900066575 |
| NK4A214_group |
| Cloacibacillus |
| Frisingicoccus |
| Shuttleworthia |
| DTU089 |
| Paludicola |
| Peptococcus |

| |
|---|
| Family_XIII_AD3011_group |
| Lachnospiraceae_ND3007_group |
| Phocea |
| Harryflintia |
| Hespellia |
| Collinsella |
| Eubacterium_nodatum_group |
| Hungatella |
| Eubacterium_xylanophilum_group |
| Lachnospiraceae_NC2004_group |
| Tyzzerella |
| Coprococcus |
| Lachnospiraceae_UCG-009 |
| UCG-009 |
| Lachnospiraceae_FCS020_group |
| UCG-005 |
| Ruminiclostridium |
| Lachnospiraceae_UCG-001 |
| Erysipelotrichaceae_UCG-003 |
| Adlercreutzia |
| Oxalobacter |
| Holdemania |
| Sanguibacteroides |
| Faecalitalea |
| Candidatus_Soleaferrea |
| Senegalimassilia |
| TM7x |

| |
|---|
| Angelakisella |
| Marvinbryantia |
| UCG-007 |
| Rothia |

FCCC partner prevalence

| Before chemotherapy | |
|---|---|
| *Genus, supplementary information >1%* | |
| Prevotella_7 | 2.98% |
| Prevotella_9 | 2.43% |
| Ruminococcus | 2.08% |
| Prevotellaceae_Ga6A1_group | 1.81% |
| Parabacteroides | 1.65% |
| Incertae_Sedis | 1.24% |
| Rikenellaceae_RC9_gut_group | 1.13% |
| Agathobacter | 1.03% |
| UCG-002 | 1.01% |

| During chemotherapy | |
|---|---|
| *Genus, supplementary information >1%* | |
| [Eubacterium]_siraeum_group | 4.29% |
| Prevotella_9 | 1.93% |
| Parabacteroides | 1.82% |
| Negativibacillus | 1.67% |
| Fusicatenibacter | 1.62% |
| Rikenellaceae_RC9_gut_group | 1.39% |
| Parasutterella | 1.29% |
| Lachnoclostridium | 1.22% |
| Prevotella_7 | 1.19% |
| Agathobacter | 1.12% |
| Barnesiella | 1.08% |
| Prevotellaceae_Ga6A1_group | 1.07% |
| Bilophila | 1.00% |

| After chemotherapey | |
|---|---|
| *Genus, supplementary information >1%* | |
| Roseburia | 3.00% |
| Agathobacter | 2.13% |
| [Eubacterium]_siraeum_group | 1.78% |
| [Eubacterium]_hallii_group | 1.72% |
| UCG-002 | 1.47% |
| Parabacteroides | 1.35% |
| Prevotellaceae_Ga6A1_group | 1.32% |
| Fusicatenibacter | 1.15% |
| Paraprevotella | 1.10% |
| Negativibacillus | 1.09% |

FCCC throughout chemotherapy Venn Supplementary

| |
|---|
| N=99 |
| Bacteroides |
| Faecalibacterium |
| Alistipes |
| Blautia |
| Christensenellaceae_R-7_group |
| UCG-002 |
| Phascolarctobacterium |
| [Eubacterium]_siraeum_group |
| Lachnoclostridium |
| [Ruminococcus]_torques_group |
| NK4A214_group |
| Negativibacillus |
| Oscillibacter |
| UCG-005 |
| Sutterella |
| Parabacteroides |
| [Eubacterium]_hallii_group |
| Anaeroplasma |
| Intestinimonas |
| Colidextribacter |
| UCG-003 |
| Agathobacter |
| Ruminococcus |
| Coprococcus |
| Subdoligranulum |
| Klebsiella |
| Parasutterella |

| |
|---|
| Fusicatenibacter |
| Anaerostipes |
| Oscillospira |
| Flavonifractor |
| Prevotella |
| Alloprevotella |
| Dorea |
| Lachnospira |
| Dialister |
| Comamonas |
| Tuzzerella |
| Odoribacter |
| Cloacibacillus |
| [Eubacterium]_ruminantium_group |
| Desulfovibrio |
| Lachnospiraceae_ND3007_group |
| Family_XIII_AD3011_group |
| [Ruminococcus]_gauvreauii_group |
| [Ruminococcus]_gnavus_group |
| Monoglobus |
| Bilophila |
| Ruminiclostridium |
| Butyricimonas |
| CAG-56 |
| Shuttleworthia |
| Candidatus_Soleaferrea |
| UBA1819 |
| Eisenbergiella |

| |
|---|
| Roseburia |
| Anaerotruncus |
| Clostridium_sensu_stricto_1 |
| Frisingicoccus |
| DTU089 |
| Collinsella |
| Victivallis |
| Butyricicoccus |
| Romboutsia |
| UCG-009 |
| Lachnospiraceae_FCS020_group |
| Lachnospiraceae_NK4B4_group |
| Marvinbryantia |
| Lachnospiraceae_AC2044_group |
| Oxalobacter |
| Hungatella |
| Erysipelotrichaceae_UCG-003 |
| Adlercreutzia |
| Defluviitaleaceae_UCG-011 |
| Intestinibacter |
| Paludicola |
| Barnesiella |
| Sellimonas |
| Holdemania |
| Pseudoflavonifractor |
| Peptococcus |
| Catenibacterium |
| Anaerofilum |

| |
|---|
| [Eubacterium]_oxidoreducens_group |
| Robinsoniella |
| UCG-004 |
| Phocea |
| Coprobacter |
| GCA-900066575 |
| Lachnospiraceae_UCG-010 |
| Erysipelatoclostridium |
| UC5-1-2E3 |
| Papillibacter |
| UCG-007 |
| Family_XIII_UCG-001 |
| Streptococcus |
| Porphyromonas |
| [Eubacterium]_ventriosum_group |
| [Eubacterium]_xylanophilum_group |

Murine model overall characteristics water vs BBN venn diagram

| | |
|---|---|
| N=44 | Lachnospiraceae_UCG-001 |
| A2 | Lachnospiraceae_UCG-006 |
| Akkermansia | Lactobacillus |
| Alistipes | Ligilactobacillus |
| Anaerofustis | Marvinbryantia |
| Anaeroplasma | Monoglobus |
| Bacteroides | Moryella |
| Bifidobacterium | Oscillibacter |
| Butyricicoccus | Paludicola |
| Candidatus_Saccharimonas | Papillibacter |
| Clostridium innocuum group | Parvibacter |
| Colidextribacter | Romboutsia |
| Coriobacteriaceae_UCG-002 | Roseburia |
| Desulfovibrio | Ruminococcus |
| Enterorhabdus | Ruminococcus_gnavus_group |
| Erysipelatoclostridium | Turicibacter |
| Eubacterium_brachy_group | Tuzzerella |
| Eubacterium_fissicatena_group | |
| Eubacterium_nodatum_group | |
| Eubacterium_ventriosum_group | |
| Eubacterium_xylanophilum_group | |
| Faecalibaculum | |
| GCA-900066575 | |
| HT002 | |
| Incertae_Sedis | |
| Intestinimonas | |
| Lachnoclostridium | |
| Lachnospiraceae_NK4A136_group | |

Murine Model Venn Diagram Supplementary

| N=45 |
| --- |
| A2 |
| Akkermansia |
| Alistipes |
| Anaerofustis |
| Anaeroplasma |
| Anaerovorax |
| Bacteroides |
| Bifidobacterium |
| Butyricicoccus |
| Candidatus_Saccharimonas |
| Colidextribacter |
| Coriobacteriaceae_UCG-002 |
| Desulfovibrio |
| Enterococcus |
| Enterorhabdus |
| Erysipelatoclostridium |
| Eubacterium_brachy_group |
| Eubacterium_fissicatena_group |
| Eubacterium_nodatum_group |
| Eubacterium_ventriosum_group |
| Eubacterium_xylanophilum_group |
| Faecalibaculum |
| Family_XIII_AD3011_group |
| GCA-900066575 |
| HT002 |
| Incertae_Sedis |
| Intestinimonas |
| Lachnoclostridium |
| Lachnospiraceae_NK4A136_group |
| Lachnospiraceae_UCG-001 |
| Lachnospiraceae_UCG-006 |
| Lactobacillus |
| Ligilactobacillus |
| Marvinbryantia |
| Monoglobus |
| Moryella |
| Oscillibacter |
| Paludicola |
| Papillibacter |
| Parvibacter |
| Romboutsia |
| Ruminococcus |
| Ruminococcus_gnavus_group |
| Turicibacter |
| Tuzzerella |

CW Group Community Venn Diagram

| |
|---|
| **N=40** |
| Faecalibacterium |
| UCG-002 |
| Subdoligranulum |
| Alistipes |
| Parabacteroides |
| CAG-352 |
| Roseburia |
| Desulfovibrio |
| Dorea |
| Odoribacter |
| Clostridium_sensu_stricto_1 |
| UCG-003 |
| CAG-56 |
| Lachnospira |
| Colidextribacter |
| Bacteroides |
| Sutterella |
| Lachnospiraceae_UCG-004 |
| Butyricimonas |
| Ruminococcus |
| Romboutsia |
| Flavonifractor |
| Monoglobus |
| Lachnoclostridium |
| Paludicola |
| Butyricicoccus |
| Ruminococcus_gauvreauii_group |

| |
|---|
| Bilophila |
| Peptococcus |
| Lachnospiraceae_ND3007_group |
| Oxalobacter |
| Intestinimonas |
| DTU089 |
| Anaerotruncus |
| Adlercreutzia |
| Erysipelotrichaceae_UCG-003 |
| Eubacterium_eligens_group |
| Harryflintia |
| Angelakisella |
| Lachnospiraceae_UCG-010 |
| **N=30** |
| Klebsiella |
| Phascolarctobacterium |
| Prevotella_7 |
| Negativibacillus |
| Eubacterium_ruminantium_group |
| Blautia |
| Eubacterium_hallii_group |
| Intestinibacter |
| Oscillospira |
| Fusicatenibacter |
| Ruminococcus_torques_group |
| Family_XIII_AD3011_group |
| Coprobacter |

| |
|---|
| UCG-005 |
| Oscillibacter |
| Hespellia |
| NK4A214_group |
| GCA-900066575 |
| Sanguibacteroides |
| Tuzzerella |
| Phocea |
| Lachnospiraceae_FCS020_group |
| Collinsella |
| Anaerofilum |
| Caproiciproducens |
| UCG-009 |
| Porphyromonas |
| Sellimonas |
| Cloacibacillus |
| Actinomyces |

FCCC cohort Venn Diagram during chemotherapy

| |
|---|
| N=102 |
| [Eubacterium]_eligens_group |
| [Eubacterium]_hallii_group |
| [Eubacterium]_nodatum_group |
| [Eubacterium]_ruminantium_group |
| [Eubacterium]_siraeum_group |
| [Eubacterium]_xylanophilum_group |
| [Ruminococcus]_gauvreauii_group |
| [Ruminococcus]_gnavus_group |
| [Ruminococcus]_torques_group |
| Adlercreutzia |
| Agathobacter |
| Alistipes |
| Anaerofilum |
| Anaerostipes |
| Anaerotruncus |
| Angelakisella |
| Bacteroides |
| Barnesiella |
| Bilophila |
| Blautia |
| Butyricicoccus |
| Butyricimonas |
| CAG-352 |
| CAG-56 |
| Candidatus_Soleaferrea |
| Candidatus_Stoquefichus |
| Caproiciproducens |

| |
|---|
| Catenibacillus |
| Cloacibacillus |
| Clostridium_sensu_stricto_1 |
| Colidextribacter |
| Collinsella |
| Coprobacter |
| Desulfovibrio |
| Dorea |
| DTU089 |
| Eisenbergiella |
| Erysipelatoclostridium |
| Erysipelotrichaceae_UCG-003 |
| Escherichia-Shigella |
| Faecalibacterium |
| Faecalitalea |
| Family_XIII_AD3011_group |
| FD2005 |
| Flavonifractor |
| Frisingicoccus |
| Fusicatenibacter |
| GCA-900066575 |
| Harryflintia |
| Hespellia |
| Holdemania |
| Howardella |
| Hungatella |
| Intestinibacter |
| Intestinimonas |

| |
|---|
| Klebsiella |
| Lachnoclostridium |
| Lachnospira |
| Lachnospiraceae_FCS020_group |
| Lachnospiraceae_NC2004_group |
| Lachnospiraceae_ND3007_group |
| Lachnospiraceae_UCG-001 |
| Lachnospiraceae_UCG-004 |
| Lachnospiraceae_UCG-010 |
| Marvinbryantia |
| Monoglobus |
| Negativibacillus |
| Odoribacter |
| Oscillibacter |
| Oscillospira |
| Oxalobacter |
| Paludicola |
| Parabacteroides |
| Paraprevotella |
| Parasutterella |
| Peptococcus |
| Phascolarctobacterium |
| Phocea |
| Prevotella |
| Prevotella_7 |
| Prevotella_9 |
| Prevotellaceae_Ga6A1_group |
| Pseudoflavonifractor |

| |
|---|
| Rikenellaceae_RC9_gut_group |
| Robinsoniella |
| Romboutsia |
| Roseburia |
| Ruminiclostridium |
| Ruminococcus |
| Sanguibacteroides |
| Sellimonas |
| Senegalimassilia |
| Shuttleworthia |
| Subdoligranulum |
| Sutterella |
| TM7x |
| Tyzzerella |
| UBA1819 |
| UC5-1-2E3 |
| UCG-002 |
| UCG-003 |
| Veillonella |

FCCC Group Community Venn Diagram

| |
|---|
| **N=40** |
| Faecalibacterium |
| UCG-002 |
| Subdoligranulum |
| Alistipes |
| Parabacteroides |
| CAG-352 |
| Roseburia |
| Desulfovibrio |
| Dorea |
| Odoribacter |
| Clostridium_sensu_stricto_1 |
| UCG-003 |
| CAG-56 |
| Lachnospira |
| Colidextribacter |
| Bacteroides |
| Sutterella |
| Lachnospiraceae_UCG-004 |
| Butyricimonas |
| Ruminococcus |
| Romboutsia |
| Flavonifractor |
| Monoglobus |
| Lachnoclostridium |
| Paludicola |
| Butyricicoccus |
| Ruminococcus_gauvreauii_group |

| |
|---|
| Bilophila |
| Peptococcus |
| Lachnospiraceae_ND3007_group |
| Oxalobacter |
| Intestinimonas |
| DTU089 |
| Anaerotruncus |
| Adlercreutzia |
| Erysipelotrichaceae_UCG-003 |
| Eubacterium_eligens_group |
| Harryflintia |
| Angelakisella |
| Lachnospiraceae_UCG-010 |
| **N=30** |
| Klebsiella |
| Phascolarctobacterium |
| Prevotella_7 |
| Negativibacillus |
| Eubacterium_ruminantium_group |
| Blautia |
| Eubacterium_hallii_group |
| Intestinibacter |
| Oscillospira |
| Fusicatenibacter |
| Ruminococcus_torques_group |
| Family_XIII_AD3011_group |
| Coprobacter |

| |
|---|
| UCG-005 |
| Oscillibacter |
| Hespellia |
| NK4A214_group |
| GCA-900066575 |
| Sanguibacteroides |
| Tuzzerella |
| Phocea |
| Lachnospiraceae_FCS020_group |
| Collinsella |
| Anaerofilum |
| Caproiciproducens |
| UCG-009 |
| Porphyromonas |
| Sellimonas |
| Cloacibacillus |
| Actinomyces |

| | |
|---|---|
| P1a | |
| Prevotella | 2.54% |
| Clostridium_sensu_stricto_1 | 1.28% |
| P1b | |
| Prevotella | 5.03% |
| Eubacterium_xylanophilum_group | 2.19% |
| Lachnospiraceae_UCG-001 | 1.13% |
| P1c | |
| UCG-003 | 1.37% |
| Lachnospiraceae_UCG-001 | 1.22% |
| Eubacterium_xylanophilum_group | 1.20% |
| P2a | |
| Haemophilus | 2.26% |
| Streptococcus | 2.00% |
| Colidextribacter | 1.55% |
| Ruminococcus_torques_group | 1.47% |
| Bilophila | 1.06% |
| P2b | |
| Bilophila | 2.22% |
| Butyricimonas | 1.10% |
| P2c | |
| - | |
| P3a | |
| Desulfovibrio | 1.99% |
| Dorea | 1.27% |
| P3b | |
| UCG-003 | 1.60% |
| Dorea | 1.25% |
| Clostridium_sensu_stricto_1 | 1.25% |
| Desulfovibrio | 1.15% |
| Colidextribacter | 1.13% |
| P3c | |
| Eubacterium_ruminantium_group | 3.87% |
| Christensenellaceae_R-7_group | 1.07% |
| P4a | |
| Lachnospiraceae_UCG-004 | 1.77% |
| Anaerostipes | 1.77% |
| Ruminococcus_gauvreauii_group | 1.54% |
| Romboutsia | 1.31% |
| P4b | |
| Ruminococcus_gauvreauii_group | 2.36% |
| Tuzzerella | 1.97% |
| Ruminococcus_torques_group | 1.09% |
| P4c | |
| - | |
| P5a | |
| - | |
| P5c | |
| Romboutsia | 1.25% |
| P6a | |
| UCG-005 | 4.59% |
| Eubacterium_ruminantium_group | 3.92% |
| Lachnospiraceae_UCG-003 | 3.45% |
| NK4A214_group | 1.88% |
| Desulfovibrio | 1.25% |
| Lachnospiraceae_UCG-004 | 1.16% |
| Eubacterium_xylanophilum_group | 1.04% |
| P7a | |
| Sutterella | 2.03% |
| Oscillibacter | 1.33% |
| Flavonifractor | 1.31% |
| Coprobacter | 1.25% |
| P7b | |
| Oscillibacter | 1.80% |
| Sutterella | 1.46% |
| P7c | |
| Sutterella | 1.57% |

FCC NAC response Venn Diagram

| |
|---|
| N=86 |
| Faecalibacterium |
| Bacteroides |
| Phascolarctobacterium |
| Blautia |
| Subdoligranulum |
| Alistipes |
| Parabacteroides |
| Eubacterium_hallii_group |
| Roseburia |
| Lachnoclostridium |
| Negativibacillus |
| UCG-002 |
| Eubacterium_siraeum_group |
| Paraprevotella |
| Agathobacter |
| Fusicatenibacter |
| Barnesiella |
| CAG-352 |
| Odoribacter |
| Eubacterium_eligens_group |
| Colidextribacter |
| Ruminococcus_gauvreauii_group |
| Flavonifractor |
| Sutterella |
| Lachnospira |
| Oscillibacter |
| Bilophila |

| |
|---|
| Dorea |
| Oscillospira |
| CAG-56 |
| Ruminococcus_torques_group |
| Eubacterium_ruminantium_group |
| Butyricimonas |
| Desulfovibrio |
| Lachnospiraceae_UCG-004 |
| Anaerostipes |
| Romboutsia |
| UCG-003 |
| Coprobacter |
| Clostridium_sensu_stricto_1 |
| Ruminococcus |
| Butyricicoccus |
| Tuzzerella |
| Intestinimonas |
| Monoglobus |
| Anaerotruncus |
| Lachnospiraceae_UCG-010 |
| GCA-900066575 |
| NK4A214_group |
| Cloacibacillus |
| Frisingicoccus |
| Shuttleworthia |
| DTU089 |
| Paludicola |
| Peptococcus |

| |
|---|
| Family_XIII_AD3011_group |
| Lachnospiraceae_ND3007_group |
| Phocea |
| Harryflintia |
| Hespellia |
| Collinsella |
| Eubacterium_nodatum_group |
| Hungatella |
| Eubacterium_xylanophilum_group |
| Lachnospiraceae_NC2004_group |
| Tyzzerella |
| Coprococcus |
| Lachnospiraceae_UCG-009 |
| UCG-009 |
| Lachnospiraceae_FCS020_group |
| UCG-005 |
| Ruminiclostridium |
| Lachnospiraceae_UCG-001 |
| Erysipelotrichaceae_UCG-003 |
| Adlercreutzia |
| Oxalobacter |
| Holdemania |
| Sanguibacteroides |
| Faecalitalea |
| Candidatus_Soleaferrea |
| Senegalimassilia |
| TM7x |

| |
|---|
| Angelakisella |
| Marvinbryantia |
| UCG-007 |
| Rothia |
